# The FDA-approved drug apremilast suppresses alcohol intake: clinical and pre-clinical validation

**DOI:** 10.1101/2021.05.13.444033

**Authors:** Kolter B. Grigsby, Regina A. Mangieri, Amanda J. Roberts, Marcelo F. Lopez, Alexander Tran, Evan J. Firsick, Kayla G. Townsley, Alan Beneze, Jessica Bess, Toby K. Eisenstein, Joseph J. Meissler, John M. Light, Jenny Miller, Susan Quello, Farhad Shadan, Michael Skinner, Heather C. Aziz, Pamela Metten, Richard A. Morissett, John C. Crabbe, Marisa Roberto, Howard C. Becker, Barbara J. Mason, Angela R. Ozburn

## Abstract

Treatment options for Alcohol Use Disorders (AUD) have minimally advanced since 2004, while the annual deaths and economic toll have become alarmingly high. Bringing potential therapeutics beyond the bench and into the clinic for AUD requires rigorous pharmacological screening across molecular, behavioral, pre-clinical, and clinical studies in neuroscience. The repurposing of FDA-approved compounds is an effective and expedited means of screening pharmacotherapies for AUD. Here, we demonstrate that apremilast, a phosphodiesterase type 4 inhibitor that is FDA approved for psoriasis and psoriatic arthritis, reduces binge-like alcohol intake and behavioral measures of motivation in unique, preclinical genetic risk models for drinking to intoxication and reduces excessive alcohol drinking in models of stress-facilitated drinking and alcohol dependence. In a double blind, placebo-controlled human laboratory study in non-treatment seeking individuals with AUD, apremilast significantly reduced the number of drinks per day. Lastly, using site-directed drug infusions and electrophysiology we determined that apremilast may act by increasing neural activity in the nucleus accumbens, an important alcohol-related brain region, to reduce alcohol intake in mice. These results demonstrate that apremilast reduces excessive alcohol drinking across a spectrum of AUD severity and support its importance as a potential therapeutic for AUD.

## Introduction

Alcohol Use Disorder (AUD) is a complex psychiatric disease with far reaching impacts on society, including >95,000 associated deaths annually in the United States (U.S.) and a net economic cost of $249 billion annually (or $807/U.S. individual)^1^. Despite a growing knowledge of important genetic and molecular mechanisms, pharmacological treatment options for AUD have not advanced since the U.S. Food and Drug Administration (FDA) approval of acamprosate in 2004^2,3^. Substantial work supports immune and inflammatory pathways as critical regulators of AUDs at all stages of the disease; namely binge drinking, increased motivation to drink, and alcohol dependence^4–6^. In particular, the cyclic adenosine monophosphate (cAMP)-specific phosphodiesterase, PDE4, has gained recent attention as a potential molecular target for treating AUD^7,8^.

For every clinical drug that reaches FDA approval, there are over a thousand that failed^9^. This remains true for AUD treatment options, whereby numerous promising preclinical drugs have proven ineffective in clinical trials^2^. To advance the translational efficacy of preclinical alcohol research there needs to be an emphasis on re-purposing currently FDA approved compounds as well as a collaborative effort to test promising pharmacotherapies across meaningful drinking paradigms, strains, and species^10^. Moreover, such studies can help inform and support clinical trials.

A central goal of the present study was to determine whether the currently FDA-approved PDE4 inhibitor, apremilast, may reduce ethanol drinking across the progression of AUD by testing its efficacy in relevant preclinical drinking paradigms, genetic animal models, and human subjects. Specifically, the effects of apremilast were evaluated in 5 clinically relevant animal models of excessive alcohol drinking (listed in order of increasing chronicity): 1) binge-like drinking^11^, 2) motivation for self-administration^12^, 3) drinking despite negative consequences (a model of compulsive-like alcohol drinking)^13–15^, 4) stress-facilitated escalation of drinking^16^, and 5) dependence-induced escalation of drinking^17,18^. To complement and extend our preclinical behavioral genetics and pharmacology studies, a double-blind, placebo controlled clinical proof-of-concept study was conducted to determine the effects of apremilast in non-treatment seeking individuals with AUD.

We further studied the role of PDE4 in the nucleus accumbens (NAc) on drinking behavior and physiology using unique genetic mouse models. An extensive body of literature supports the NAc as a critical regulator of alcohol drinking^19–24^. Therefore, we sought to determine 1) whether administration of apremilast into the nucleus accumbens would be sufficient to reduce binge-like drinking and achieved blood alcohol levels and 2) whether apremilast differentially alters physiology in two types of medium spiny neurons (MSNs; dopamine receptor D1 or D2 expressing MSNs), which comprise the two major output pathways from the accumbens. Taken together, these studies provide an integrative and rigorous framework supporting further testing of the importance of apremilast as a pharmacotherapy in the treatment of AUDs.

## Results

### Apremilast reduces binge-like drinking behavior in mice selectively bred for drinking to intoxication

To test whether PDE4 inhibition reduces binge-like alcohol drinking, we administered apremilast to selectively bred “High Drinking in the Dark” (HDID-1, HDID-2) mice of both sexes prior to measuring limited access drinking using the widely adopted “Drinking in the Dark” assay^11^ (DID). HDID mice were selectively bred for drinking to intoxication as measured by blood alcohol levels (BALs) achieved during the DID task. HDID mice reliably reach BALs over 200 mg% (> 80 mg% is considered intoxicating^25^). Importantly, the HDID-1 and -2 lines were independently selectively bred from genetically heterogeneous stock (HS/Npt), and thus represent two unique genetic models of high risk for binge drinking behavior. Here we found that two clinically relevant doses of apremilast, 20 and 40 mg/kg, i.p., reduced binge drinking, as well as BALs, in female and male HDID-1 mice (Fig. 1 a,b). The 40 mg/kg dose reduced BALs to a level that is near the NIAAA definition of intoxication (80 mg%), whereas mice in the 0 mg/kg dose group achieved BALs that are nearly 2.5 times greater than the threshold for intoxication. The same doses of apremilast reduced binge-like drinking and BALs in female and male HDID-2 mice (Fig. 1 c,d). Although selected for the same phenotype, HDID-1 and HDID-2 mice display key differences in brain-related gene co-expression patterns (with HDID-2 sharing closer genetic similarity to C57BL/6J mice; another high drinking strain^26^). These findings highlight the ability of apremilast to reduce drinking to intoxication in two unique genetic animal models of risk for excessive alcohol drinking.

**Figure 1:**
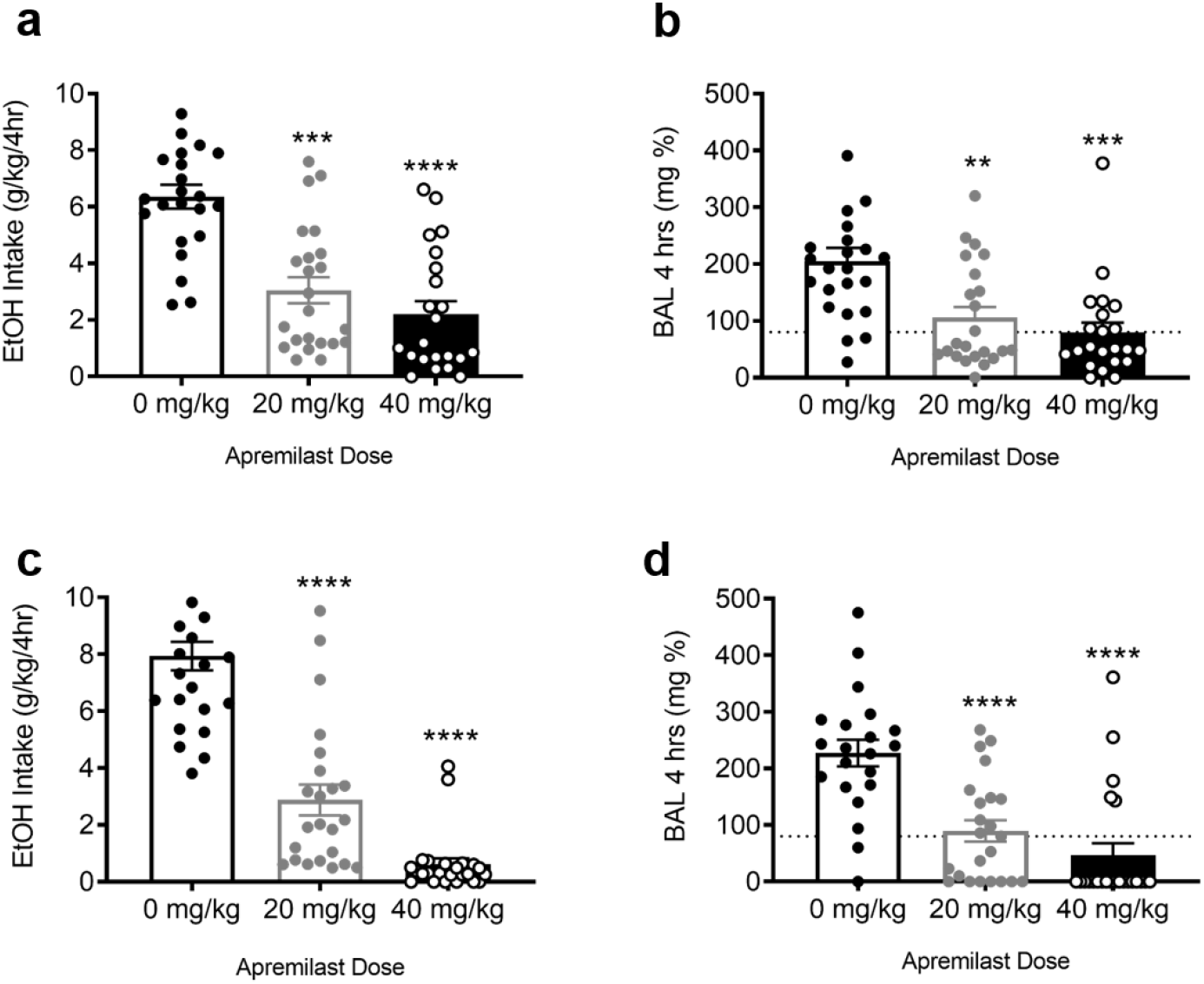
Apremilast reduces binge-like drinking behavior in mice selectively bred for drinking to intoxication. **a**, Binge-like ethanol intake (g/kg/4hrs) for HDID-1 (n = 10-12/sex/apremilast dose; main effect of apremilast [F(2,61) = 21.0, p<0.0001], with no sex or sex X treatment interactions; both doses of apremilast reduced ethanol intake in HDID-1 mice. **b**, Blood alcohol levels (mg %) in HDID-1; main effect of apremilast [F (2,64) = 9.73, p < 0.001]; both doses of apremilast reduced BALs compared to 0 mg/kg. **c**, Binge-like ethanol intake (g/kg/4hrs) for HDID-2 (n = 11-12/sex/apremilast dose); main effect of apremilast [F(2,68) = 73.2, p<0.0001]; both doses of apremilast reduced ethanol intake in HDID-2 mice. **d**, Blood alcohol levels (mg %) in HDID-2; main effect of apremilast [F (2,64) = 9.73, p < 0.001]; both doses of apremilast reduced BALs compared to 0 mg/kg. (* = *p* < 0.05, ** = p < 0.005, *** = *p* < 0.001, **** = *p* < 0.0001). Dashed line indicates legal level of intoxication (80 mg %).

To better evaluate its effects on binge-drinking behavior, we next tested whether apremilast altered drinking of other fluids using the same DID test in a previously established serial testing fashion^27^. In week 2, we measured the effect of apremilast on water drinking for HDID-1 and HDID-2 mice of both sexes (Extended Data Fig. 1 a,b). To determine whether the reduction in binge-like drinking following apremilast treatment was specific to the reinforcing/rewarding value of alcohol we tested its effects on intake of a highly palatable saccharin solution during the 3rd week of testing (Extended Data Fig. 1 c,d). Here we found that apremilast had no effect on either water intake or saccharin intake in female and male HDID-1 mice, suggesting the observed reduction in ethanol intake in HDID-1 mice was not likely due to sedation, sickness, or altered sensitivity to rewarding solutions. This supports that the effect in HDID-1 mice is specific to ethanol as a reinforcer. In contrast, apremilast reduced HDID-2 water intake in week 2 of testing and saccharin intake in week 3 (Extended Data Fig. 1 b,d). This may suggest that the reductions in alcohol drinking following apremilast treatment in HDID-2 mice are due to general effects on liquid intake and/or malaise. More importantly, despite having similar reductions in ethanol intake and BALs, the apparent difference in water and saccharin intake for HDID-1 and - 2 mice treated with apremilast highlights the value of testing multiple animal strains.

To better understand the role of PDE4 in binge-like drinking, the PDE4 inhibitor, rolipram, was administered to female and male HDID-1 mice prior to DID. All three doses of rolipram tested (5, 7.5, and 10 mg/kg) reduced alcohol drinking and BALs (Extended Data Fig. 1 e,f). The ability of two different PDE4 inhibitor compounds to reduce drinking suggests that PDE4 is likely an important regulator of excessive alcohol drinking by animals with genetic risk and is consistent with prior studies of PDE4 inhibitors in other strains^28,29^.

Because AUDs are characterized by chronic excessive alcohol drinking, we next tested the efficacy of apremilast to reduce alcohol intake in the context of chronic binge drinking. Here, we found that 40 mg/kg of apremilast reduced binge-drinking in HDID-1 mice (of both sexes) over a 4-week period compared to baseline drinking levels (Extended Data Fig. 1 g). Of note, we observed an increase in drinking after treatment ended (washout drinking levels, Extended Data Fig. 1 g). This suggests that termination of apremilast may lead to an increase in binge-drinking. There was no effect of apremilast on BALs collected on the last day of alcohol drinking during washout (Extended Data Fig. 1 h).

### Apremilast reduces the motivation for alcohol drinking in mice selectively bred for drinking to intoxication

To determine whether PDE4 inhibition reduces the motivation for alcohol drinking, we next tested the effects of apremilast in inbred HDID-1 (iHDID-1) mice during two operant ethanol self-administration tasks, both of which model and test important aspects of human motivation for alcohol^12,30,31^. We first evaluated the efficacy of apremilast to reduce the number of maximum responses mice would perform to gain access to an alcohol solution under a progressive ratio (PR) schedule of reinforcement, whereby the threshold number of lever-presses needed to gain alcohol access is rapidly increased. The schedule of operant training and testing is shown in Extended Data Fig. 2 a. The highest response ratio reached (the breakpoint) within this test session is considered a reliable measure of the motivation for alcohol in mice ^12^. Here we found that a clinically relevant dose of apremilast (40 mg/kg, i.p.) reduced the motivation for alcohol in female and male iHDID-1 mice (Fig. 2 a). This dose of apremilast also reduced the total number of operant reinforcers earned during PR (Extended Data Fig.2 b), suggesting a reduction in the reinforcing efficacy of alcohol. Consistent with our earlier findings, apremilast was also effective at reducing alcohol intake during the PR test (Extended Data Fig. 2 c); however, BALs were not evaluated because these mice were subsequently tested in another important motivation-related task (Extended Data Fig. 2 a).

**Figure 2:**
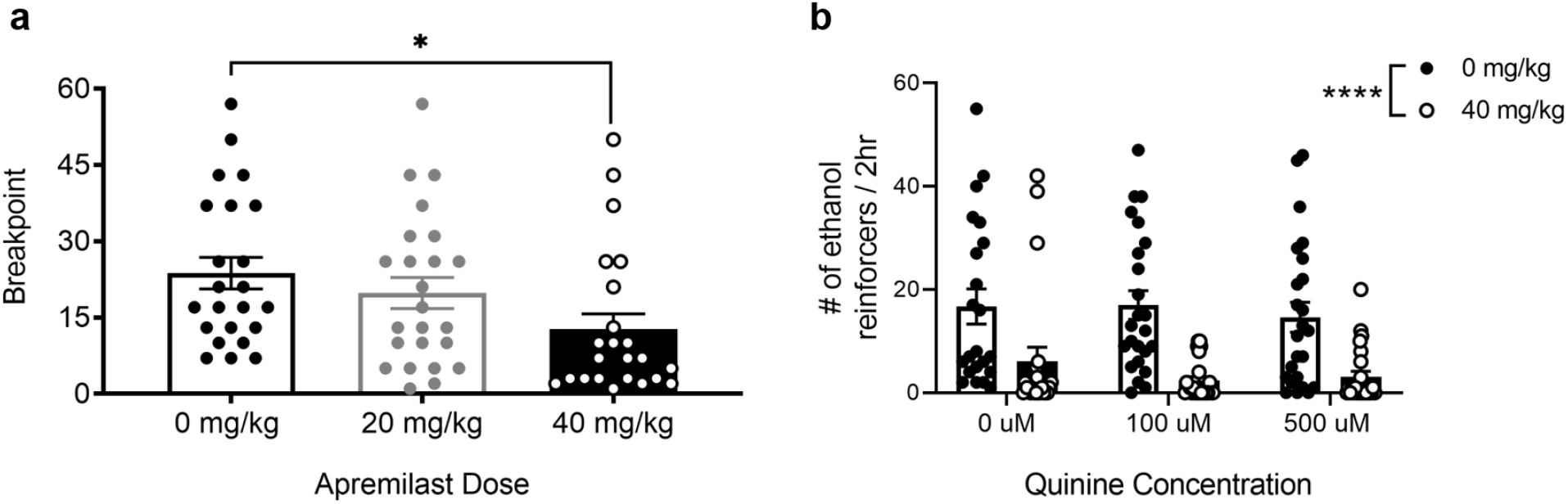
Apremilast reduces the motivation for ethanol in iHDID-1 mice. **a**, Highest operant response ratio reached (Breakpoint) during PR testing (marker of ethanol motivation) for iHDID-1 (n = 10/12/sex/apremilast treatment); main effect of treatment [F(2,64) = 4.47; p < 0.05]; 40 mg/kg reduced breakpoint iHDID-1 mice. **b**, Ethanol reinforcers earned during quinine-adulterated testing; main effect of apremilast treatment [F(1,134) = 37.90; p < 0.0001], with no effect of quinine or apremilast X quinine interaction; 40 mg/kg apremilast reduced the number of reinforcers earned for iHDID-1 mice at all quinine concentrations tested. (* = *p* < 0.05).

To ascertain whether apremilast would reduce compulsive-like responding for alcohol (another facet of human alcohol motivation), mice were then tested for quinine-adulterated alcohol responding (see timeline, Extended Data Fig. 2 a), using a widely adopted model of drinking despite negative consequences^13^. Alcohol drinking and operant behaviors were similar between iHDID-1 mice consuming alcohol alone (0 µM quinine), alcohol with low quinine concentration (100 µM), and alcohol with high quinine concentration (500 µM), indicating that iHDID drink despite negative consequences and demonstrate compulsive-like responding for alcohol. Mice were then given injections of 0 mg/kg and 40 mg/kg apremilast, prior to operant self-administration (with same alcohol +/− quinine solution), counterbalanced over two, 2-hr sessions (Extended Data Fig. 2 a). Apremilast reduced the number of alcohol access periods (reinforcers) earned and alcohol intake at all concentrations of quinine tested (Fig. 2 b and Extended Data Fig. 2 d). This suggests that apremilast reduced the motivation to drink despite negative consequences and taken together, these findings indicate that apremilast effectively reduces behavioral signs of alcohol motivation in mice bred to drink to intoxication.

### Apremilast reduces dependence-induced escalations in alcohol intake in C57BL/6J mice

To test whether apremilast reduces harmful drinking associated with alcohol dependence, two models of dependence-induced escalations in ethanol drinking were used in C57BL/6J mice, a strain that exhibits high alcohol drinking. The rationale for using this strain stems from the fact that both dependence-related drinking models were developed in C57BL/6J mice^16,18^. The experimental details and timelines are shown in Extended Data Fig. 3. In the first set of experiments, a daily stressor (Forced Swim Stress; FSS) was given in combination with chronic intermittent ethanol vapor exposure (CIE) to escalate drinking behavior (Fig. 3 a). Stress is thought to play a critical role in alcohol dependence, whereby forced swim stress prior to CIE exposure has been shown to enhance escalation and alcohol intake beyond CIE alone^16,17^. Consistent with published findings, C57BL/6J mice exposed to CIE and those given stress in combination with CIE (CIE + FSS) had higher ethanol intake than air control mice and those given FSS alone^16^ (Fig. 3 a). Here we found that 20 mg/kg of apremilast reduced alcohol intake in stressed, dependent (CIE+FSS) mice and that 40 mg/kg of apremilast effectively reduced ethanol intake in stressed and non-stressed, dependent mice (Fig. 3 b).

**Figure 3:**
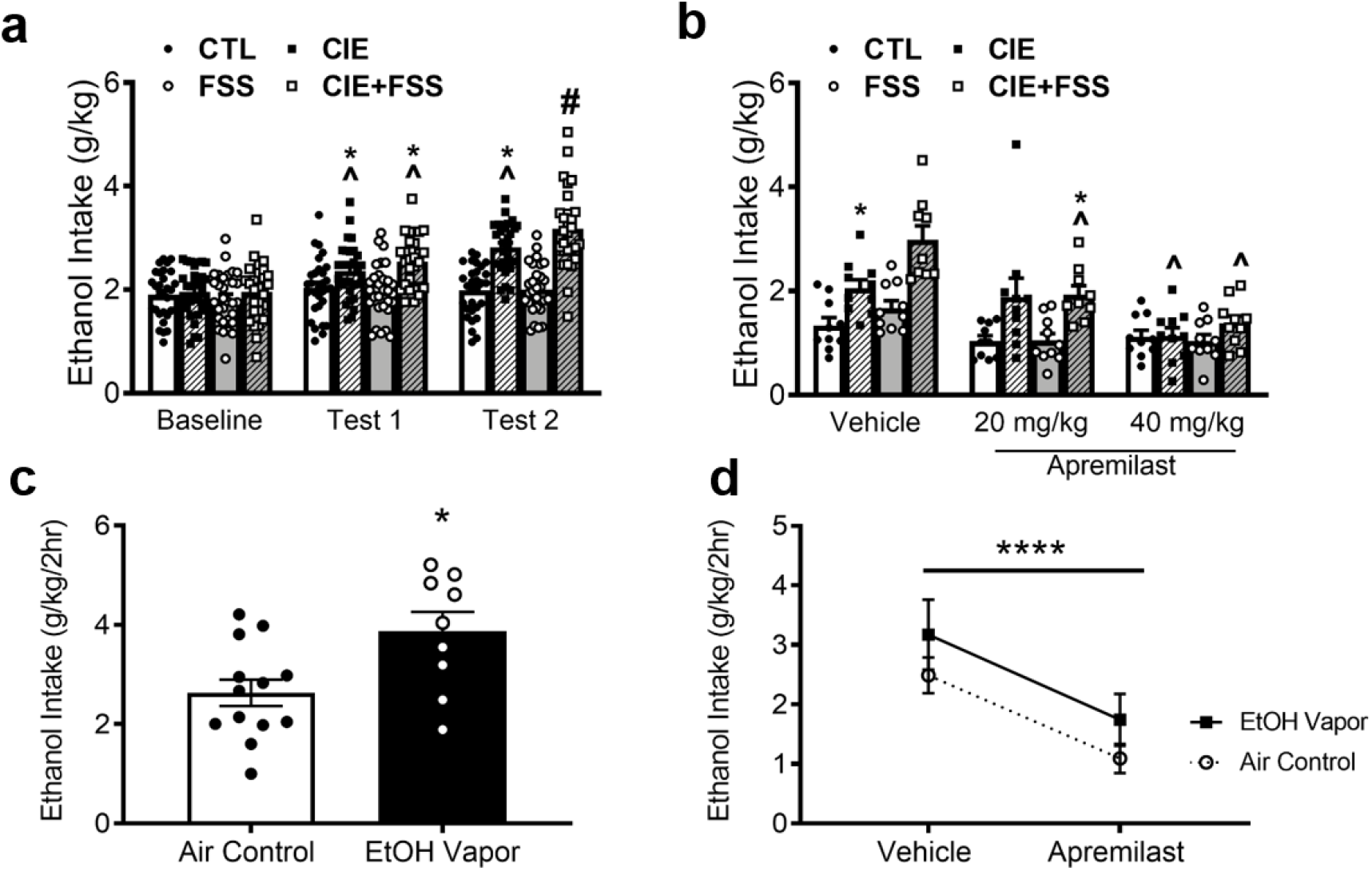
Apremilast reduces dependence induced escalations in ethanol drinking in C57BL/6J mice. **a**, Ethanol intake (g/kg/2hr) for male C57BL/6J mice (n = 9-10/vapor group/stress group/apremilast treatment) during baseline and tests 1 and 2; main effect of group [F(3,114)=15.22; p<0.001], phase [F(2,114)=60.80;p<0.001] and a group X phase interaction [F(6,228)=13.25; p<0.001]; CIE and CIE+FSS had higher intake compared to baseline and CTL values (*) and their own baseline (^); CIE+FSS had higher intake in test 2 than all other groups and their own baseline (#). **b**, Ethanol intake (g/kg/2hr) during test 3; main effect of group [F(3,106)=16.28; p<0.001], apremilast [F(2,106)=21.83;p<0.001] and a group X treatment interaction [F(6,106)=3.25; p<0.01]; for mice that received vehicle, ethanol intake was higher for CIE mice compared to CTL mice (*) and higher for CIE+FSS compared to the three groups that also received vehicle (#). CIE+FSS mice that received 20 mg/kg apremilast continued to drink more ethanol than CTL mice (*). However, this dose reduced ethanol intake compared to its vehicle condition group (^). The 40 mg/kg apremilast dose resulted in a significant decrease in ethanol intake in CIE and CIE+FSS mice compared to their vehicle equivalent (^). **c**, Ethanol intake (g/kg/2hrs) for female and male C57Bl/6J mice (n = 10/vapor group/apremilast treatment) following 3-weeks of CIE, main effect of vapor exposure, whereby ethanol vapor increased intake. **d**, Ethanol intake (g/kg/2hrs) during test week, main effect of treatment, 40 mg/kg (p.o.) reduced intake in ethanol vapor and air exposed mice (* = *p* < 0.05, *** = *p* < 0.001, **** = *p* < 0.0001).

To test the effects of apremilast in a more chronic model of dependence-induced escalations in alcohol drinking, female and male C57BL/6J mice underwent a standard CIE protocol^18,32–35^. Following 4 cycles of ethanol vapor exposure, mice showed an increase in alcohol intake relative to control mice (Fig. 3 c). When given orally prior to the last day of drinking, apremilast (20 mg/kg) was shown to decrease ethanol intake in non-dependent (air control) and dependent mice. In all, the above findings extend the efficacy of apremilast to reduce excessive ethanol drinking in two well-established animal models of alcohol dependence.

### Individuals with AUD consume less drinks per day when treated with apremilast

A Phase IIa double-blind, placebo-controlled proof-of-concept (POC) study was conducted with the aim of clinically validating the effect of apremilast on decreasing alcohol intake in preclinical models of AUD. The hypothesis being that individuals with AUD who were treated with apremilast would consume significantly fewer standard drinks (~14 grams of alcohol per drink) per day over an 11-day period of *ad libitum* drinking than those treated with placebo. A higher oral dose (90 mg/d) was used to assess efficacy for reducing drinking in individuals with AUD than the 60 mg/d dose typically prescribed for the FDA-approved indication of psoriasis. Our preclinical dose-ranging data showed doses > 20 mg/kg to be associated with decreased drinking; to facilitate translation across species, we used 90 mg as the equivalent dose in humans. Earlier PDE4 inhibitors like rolipram and ibudilast are associated with side effects, particularly nausea and vomiting, that significantly reduce patient acceptability. Apremilast shows less of the PDE4 adverse reactions clinically^36^ and has lower affinity for the PDE4D isozyme which is thought to be associated with the emetic effects of PDE4 inhibition^37^, and thus may be well tolerated in the higher than standard dosing that is indicated for reducing drinking in AUD.

Study admission criteria specified non-treatment seeking male and female paid volunteers 18 – 65 years of age with ≥ moderately severe AUD, as defined by the Diagnostic and Statistical Manual for Mental Disorders – Fifth Edition (DSM-5) criteria^38^. The *Consolidated Standards of Reporting Trials* (CONSORT) flow diagram is shown in Extended Data Figure 4. Subjects were randomly assigned in a 1:1 ratio to treatment with a target dose of 90 mg/d of apremilast or matched placebo in a parallel group design (Extended Data Table 1). The randomization code included stratification on sex and baseline C-Reactive Protein (CRP; a blood marker of inflammation) status (< 2 mg/L vs ≥ 2 mg/L) to ensure an equivalent distribution of subjects across groups on two factors potentially related to outcome. Plasma levels of cytokines (TNF-alpha, CCL2, CXXL10), cortisol, apremilast and serum endotoxin were assayed after study completion for evaluation as potential physiological moderators of treatment response (Extended Data Table 2).

The rate of study completion (84%) was equivalent across groups and is detailed in Extended Data Figure 4. Subjects were 24 (47.1%) females and 27 (52.9%) males with a mean age of 41.2 (16.3) years. Subjects had been drinking heavily for 12.3 (10.5) years and met criteria for 6.4 (2.3) DSM-5 symptoms at baseline, indicating a severe level of AUD. Apremilast (N = 26) and placebo (N = 25) groups did not differ on baseline demographic, clinical or physiological variables, as summarized in Extended Data Tables 1 and 2.

All randomized subjects (N = 51) were included in an intention-to-treat analysis that employed latent growth modeling^39^ (LGM) to compare drinking in apremilast vs. placebo groups during the 11-day period of *ad libitum* drinking (R package glmmadmb v0.8.3.3)^40^. Apremilast significantly (p = 0.025) reduced the number of drinks per day relative to placebo, as shown in Figure 4. The LGM procedure generated change values of 2.74 drinks per day for apremilast and 0.48 for placebo and yields a Cohen’s d value of 0.77 which can be interpreted as a “large” effect of apremilast on decreasing drinking^41^. No baseline demographic, clinical or physiological variable contributed significantly to drinking outcome. No serious or severe adverse events occurred. Adverse drug effects of diarrhea, nausea, abdominal pain, and somnolence were two or more times more likely with apremilast than placebo as shown in Extended Data Table 3; these effects were typically mild and were not associated with treatment discontinuation.

**Figure 4:**
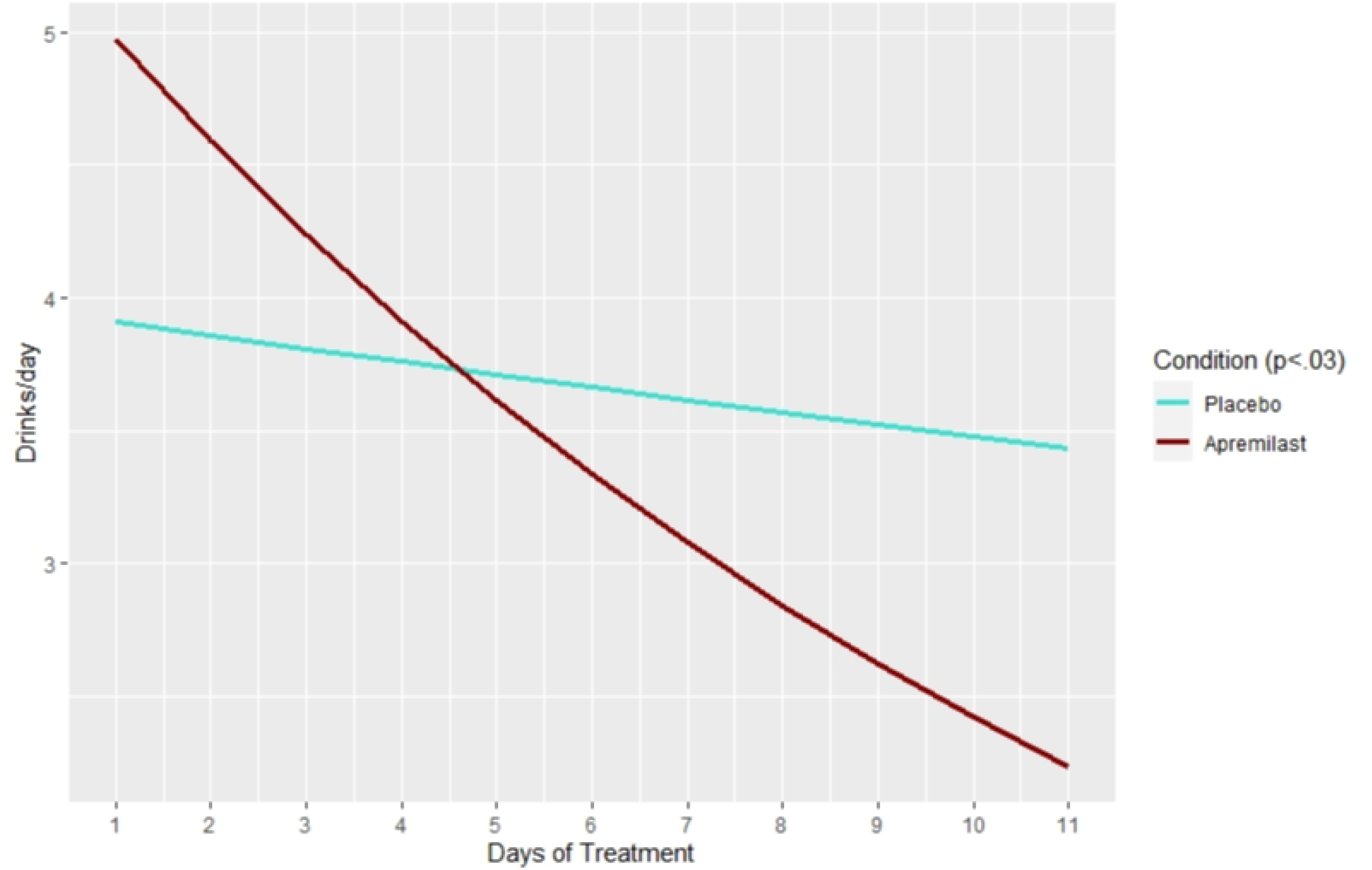
Apremilast 90mg/d significantly (p = 0.025) reduces the number of drinks per day relative to placebo in 51 non-treatment seeking individuals with alcohol use disorder of ≥ moderate severity. A latent growth curve model was used to calculate an effect size for apremilast vs. placebo in the decrease in drinks per day from baseline through 11 days of *ad libitum* drinking. This procedure generated change values of 2.74 drinks per day for apremilast and 0.48 for placebo, and yields a Cohen’s d value of 0.77, which can be interpreted as a “large” effect of apremilast on decreasing drinking.

In summary, this double-blind, placebo controlled POC study found a large effect of apremilast 90 mg/d on decreasing drinking relative to placebo in 51 non treatment seeking men and women with severe AUD. This effect size compares favorably to the modest effect sizes associated with the FDA-approved AUD drugs, acamprosate and naltrexone^42^. Results provide clinical validation of PDE4 inhibition as a general therapeutic strategy for AUD, and specifically for our extensive preclinical data showing apremilast decreases drinking in animal models of AUD. The 90 mg/d dose, while 50% higher than standard dosing for psoriasis, was well tolerated in this AUD sample. Taken together, these POC efficacy and safety data lend support to further development of apremilast as a novel treatment for AUD.

### The nucleus accumbens is a critical site of action for reduction of drinking by apremilast

Recent evidence suggests that increased expression of PDE4 subtypes, namely PDE4b, is linked to human AUD^7,8^. Therefore, we sought to determine the effects of chronic binge-like drinking on PDE4 subtype expression in the nucleus accumbens (NAc; a brain region integral to alcohol drinking) in HDID-1 mice. Here we found that binge-drinking increased the expression of both PDE4a and PDE4b (Fig. 5 a,b).

**Figure 5:**
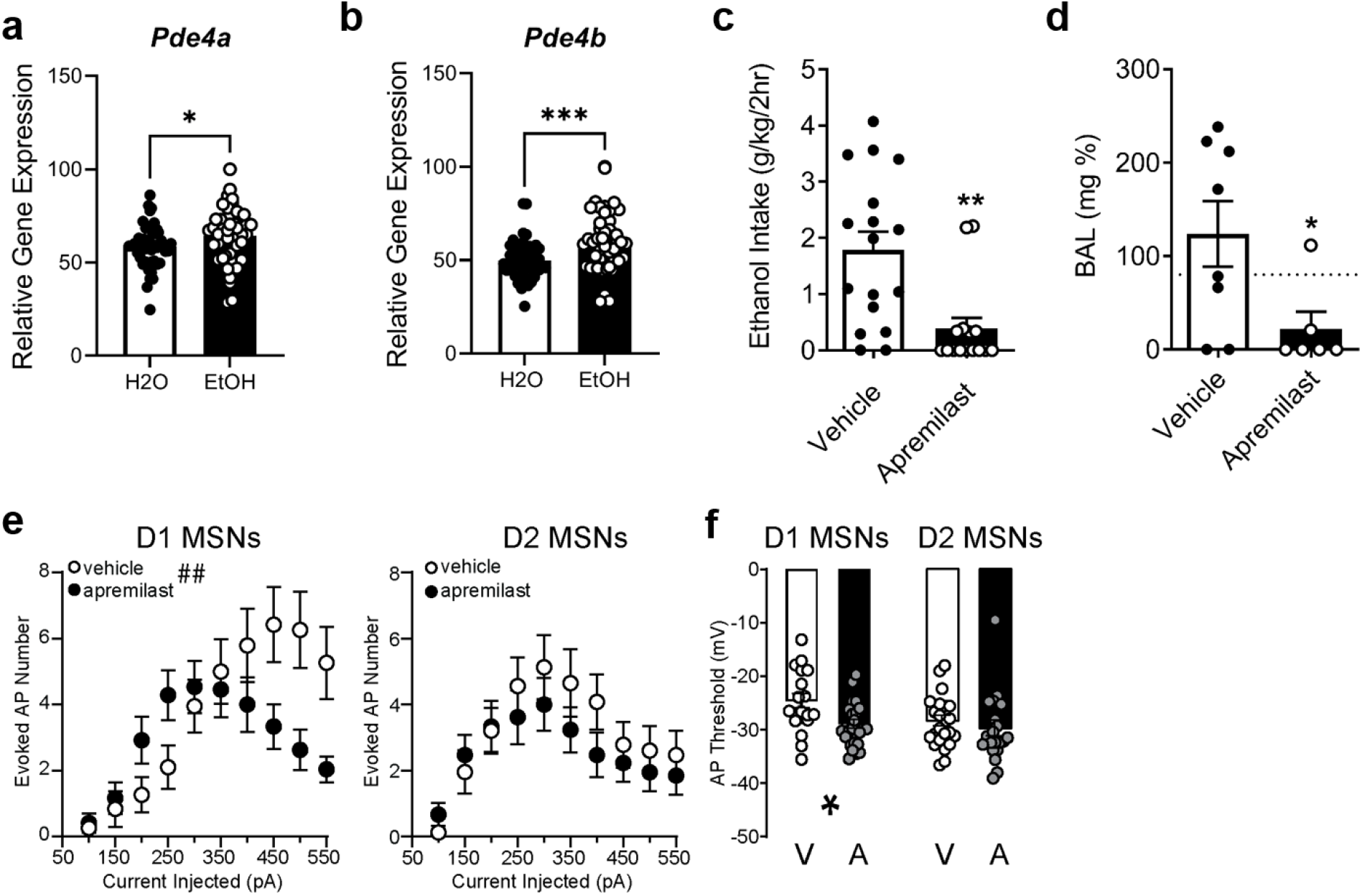
The nucleus accumbens is a critical site of action for reduction of drinking by apremilast. **a**, Relative gene expression of NAc *Pde4a* expression for female HDID-1 mice (46-48/fluid group); main effect of fluid type [F(1,79)=15.22; p<0.05] ethanol mice express higher levels of *PDE4a*. **b**, Relative gene expression of NAc *Pde4b*; main effect of fluid type [F(1,79)=13.02; p<0.05]. Ethanol mice express higher levels of *PDE4b*. **c**, Ethanol intake (g/kg/2hrs) following intra-NAc apremilast infusions (0 or 2 µg/µL/side) for male HDID-1 mice (n = 19-20/fluid group/infusion group), significant effect of apremilast (Student’s t-test; p < 0.01). **d**, Blood alcohol levels (mg %); significant effect of apremilast (Student’s t-test; p < 0.05).(* = *p* < 0.05, *** = *p* < 0.001). Dashed line indicates legal level of intoxication (80 mg %). **e**, Input-output curves showing number of action potentials (AP) evoked in response to 300 msec current steps of increasing amplitudes for D1 MSNs (n = 19 vehicle, 24 apremilast) and D2 MSNs (n = 23 vehicle, 21 apremilast). There was a 3-way interaction of Treatment X Cell Type X Current Amplitude [F(3,216)=2.89, p<0.05]. ##, Treatment X Cell Type interaction in D1 MSNs [F(2,99)=6.1, p<0.01]. **f**, Membrane potential for action potential threshold; main effect of Treatment [F(1,82) = 6.26, p < 0.05)]. * = p <0.05, effect of Treatment within D1 MSNs (Bonferroni’s multiple comparisons test). A: apremilast (1 µM), V: vehicle (0.002% DMSO). Dashed line indicates level of intoxication (80 mg %).

To determine whether inhibition of PDE4 in the NAc could reduce drinking, we next tested the effects of intra-cranial accumbens infusions of apremilast on binge drinking in HDID-1 mice. We observed a significant decrease in binge-like ethanol drinking and BALs (Fig. 5 c,d), with no effect on either water or saccharin intake (Extended Data Fig. 5).

We therefore evaluated how acute treatment with apremilast altered neuronal membrane properties and excitability in mouse NAc MSNs using *ex vivo* brain slice electrophysiology. We compared the effects of apremilast between dopamine D1 receptor-expressing (D1 MSNs) and D2 receptor-expressing MSNs (D2 MSNs), which comprise the two major output pathways of the NAc. We observed a cell type by treatment interaction effect on evoked action potential firing, with apremilast having no effect in D2 MSNs, but decreasing action potential firing induced/evoked by depolarizing currents/steps in D1 (Fig. 5 e). The apremilast effect on evoked firing was not accompanied by treatment differences in resting membrane potential, input resistance, or rheobase (Extended Data Table 4). Rather, the action potential threshold of D1 MSNs was significantly reduced by apremilast treatment (Fig. 5 f). Other action potential properties that are indicative of sustained firing capability were unaffected (Extended Table 4). Thus, our results indicate that apremilast primarily promotes D1 MSN excitability, and does so by increasing the responsiveness of these neurons to membrane depolarization.

Dopaminergic neurotransmission in NAc medium spiny neurons (MSNs) is largely mediated through PKA signaling, of which PDE4 is a critical regulator. Nishi *et al*. demonstrated that the PDE4 inhibitor, rolipram, increased neuronal excitability in isolated MSNs^43^. There is evidence demonstrating that altering activity of the NAc leads to a decrease in alcohol craving and relapse in humans^22,44,45^ and binge-like drinking in mice^23,24,46^.

Extending the importance of PDE4 inhibition to NAc-mediated ethanol drinking, the present findings show that site-specific apremilast treatment is sufficient to reduce binge-like ethanol drinking and may do so by promoting the excitability of D1 MSN outputs.

## Discussion

AUD is a complex, polygenetic disorder which requires a concerted research effort to address its underlying mechanisms. While the Human Genome Project has and will continue to be a powerful tool in understanding the genetic basis of complex traits, treatment of human disease and drug discovery still rely on the experimental rigor of behavioral pharmacology^10^. It is imperative to evaluate promising pharmacotherapies across multiple drinking paradigms, species, and strains; therefore, the present work determined whether the currently FDA approved PDE4 inhibitor, apremilast, would reduce harmful alcohol drinking in male and female mice from four different strains of mice with high genetic risk for excessive drinking (i.e. selectively bred HDID-1 and 2, inbred HDID-1 mice, and C57BL/6J mice). Strikingly, we determined that apremilast reduced harmful drinking across a spectrum of clinically relevant drinking models for binge-like, motivational, compulsive-like, and stress- and non-stress induced facilitation of dependence-like drinking.

For clinical validation of these findings, we employed a double blind, placebo-controlled study in non-treatment seeking individuals with AUD and found that oral apremilast was effective at reducing the number of daily drinks consumed. Moreover, we identified that apremilast’s effects at the level of the accumbens as important for regulating binge-like drinking and for regulating activity in a neural circuit relevant to alcohol-related behaviors^47,48^. Taken together, this collaborative set of studies from 5 independent laboratories and universities highlights apremilast as a powerful AUD treatment option and further identifies mechanisms by which apremilast may reduce harmful alcohol drinking.

Substantial evidence supports PDE4 inhibition as a likely and viable pharmacotherapeutic option for treating AUD. Others have shown that non-specific PDE inhibitors, such as ibudilast, reduce alcohol drinking in mouse and rat models of dependence^49^. Human laboratory testing revealed that ibudilast improves mood on secondary measures of stress and alcohol related cue exposures, and reduces levels of alcohol craving ^50,51^. The PDE4 inhibitor, rolipram, was first shown to reduce ethanol intake in mice given a 2-bottle choice between water and ethanol ^52^. Elegant behavioral pharmacology studies have addressed which of the PDEs are most important for regulating drinking ^8,29,53,54^, whereby Blednov *et al*. subsequently found that 9 different PDE4 inhibitors were efficacious in reducing ethanol intake and preference in a two-bottle choice, limited access test in male C57BL/6J mice^28^. Although this thorough work highlighted PDE4 inhibition as an effective means of reducing harmful drinking, blood alcohol levels were not measured and therefore PDE4’s role in reducing pharmacologically relevant BALs was not fully addressed. In a subsequent study, Ozburn *et al*. determined that the PDE4 inhibitor, rolipram, significantly reduced binge-like ethanol intake and BALs and in female and male HDID-1, HDID-2 and their heterogenous founders, the HS/Npt mice^55^. Here we show that an FDA approved PDE4 inhibitor is similarly effective at reducing binge-like ethanol drinking in the same genetic risk model of drinking to intoxication, and further ask whether apremilast is useful for reducing alcohol drinking across a spectrum of models for AUD severity.

Testing the importance of PDE4 in operant ethanol self-administration models helps to address a key component of human AUDs with face validity for clinical populations; the motivation for alcohol drinking. In 2012, Wen *et al*. demonstrated that systemic PDE4 inhibition using rolipram decreased operant responding for an alcohol (5% ethanol) but not sucrose (10%) solution in Fawn Hooded rats^56^. Although this suggests that rolipram reduced the reinforcing effects of alcohol, the motivation for or willingness of these rats to work for alcohol access was not directly tested. We assayed this directly using a progressive ratio schedule of reinforcement and found that apremilast reduced measures of alcohol motivation in iHDID-1 mice, whereby apremilast reduced the breakpoint, or effort exerted by these mice to obtain alcohol. Using a model of drinking despite negative consequences, which addresses aspects of compulsive alcohol use and alcohol motivation, we determined that apremilast also reduced responding for quinine-adulterated alcohol in iHDID-1 mice. Together, these findings suggest that apremilast is effective at reducing harmful drinking and related behaviors under the conditions of chronic use.

Investigating the role of PDE4 in models of dependence-induced escalation of drinking helps to address a key component of human AUDs with face validity for clinical populations; dependence-like alcohol drinking. The present data demonstrate that apremilast effectively reduces ethanol intake in stressed and non-stressed alcohol dependent and non-dependent mice. Notably, others have found that male C57BL/6J mice subjected to forced swim stress and CIE had persistent increases in immune-related, alcohol responsive genes, including *Pde4b* expression^57^. Moreover, previous work showed that nonspecifically targeting phosphodiesterases (via peripheral administration of ibudilast) reduced dependence-induced alcohol intake in C57BL/6J mice^49^. Together, our results highlight the therapeutic value of apremilast to reduce relapse-like alcohol drinking, a critical component of harmful drinking and AUD psychopathology. Of note, these findings were conducted at two separate institutions, which further supports the generalizability and validity of apremilast as a potential AUD treatment option.

Because apremilast works across a spectrum of models, in both sexes of four strains of mice (at multiple labs and universities) and importantly, in humans, we sought to determine the neural mechanisms by which PDE4 inhibition reduces harmful drinking. The nucleus accumbens (NAc) is an integral region for reward-related behaviors and is well studied for its critical role in ethanol-drinking, whereby structural and molecular changes following both acute and chronic ethanol drinking are thought to play a role in further aberrant drinking patterns^58^. Evidence demonstrates that electrical stimulation of the NAc leads to a decrease in alcohol craving and relapse in humans^22,44,45^ and alcohol drinking in rodents^59–61^. The findings herein show that chronic binge drinking results in increased NAc expression of two *Pde4* subtypes, *Pde4a* and *Pde4b*. Notably, heightened expression of the *Pde4b* isoform has been genetically associated with chronic ethanol intake in humans^7,8^. PDE4 inhibition has been shown to increase pre- and post-synaptic cAMP-driven markers of neuronal excitability in the NAc^43^. While this pathway is considered a regulator of harmful drinking behaviors, PDE’s are known to have complex intracellular interactions which may lead to altered neural function and changes in behavior^62^. The present findings show that site-specific apremilast treatment is sufficient to reduce binge-like alcohol drinking, demonstrating the importance of PDE4 inhibition to NAc-mediated drinking. The extent to which PDE4 inhibition, and in particular apremilast, alter the excitability of subpopulations of MSNs in the NAc helps to identify potential critical neurobiological mechanisms and may in part explain the observed reduction in harmful alcohol drinking across drinking models. Here, we saw that apremilast increased neuronal excitability in D1, but not D2 MSNs. Chemogenetic activation of these distinct populations suggests that D1 MSNs are more important for alcohol drinking^60^; therefore, our findings suggest that apremilast may be acting primarily through D1 MSNs to reduce alcohol drinking and alcohol-related behaviors.

## Acknowledgements

Supported by the NIH grants (AA016651, AA013519, AA010760, AA07468, and AA027692, U01 AA013498, DA013429); the US Department of Veterans Affairs Grants (BX000313, BX004699, and IK2 BX002488), and a gift from the John R. Andrews Family. The authors thank Dr. R. Adron Harris for his contributions during the preparation of this paper and as Director of the Integrative Neuroscience Initiative on Alcoholism – Neuroimmune consortium. We also thank Dr. Esther Maier for developing the methodology and conducting the determination of apremilast concentration in human plasma. The authors thank the following individuals for their technical assistance: Stephanie Timko, Jason Schlumbohm, Wyatt Hack, Nathan Ashby, Sarah E. Reasons.

## Contributions

**Preclinical:** ARO, KBG, RAM, AJR, MFL conceived the experiments, performed the experiments, performed analyses, and wrote the paper. EJF, AT, KGT, and HCA performed the experiments. PM analyzed the data. JCC, MR, and HCB conceived the experiments and edited the paper. **Clinical:** BJM conceived and conducted the clinical study and wrote the paper. JML analyzed the data, interpreted results and wrote the paper. AB, JB, JM, SQ, FS and MS conducted the study. TKE and JJM analyzed the physiological data and interpreted results.

## Ethics Declaration

Competing Interests: The authors declare no competing interests.

## Data Availability

All data are available upon request, without restriction.

## Extended Data

**Extended Data Table 1.** Baseline demographic and clinical characteristics of apremilast and placebo groups (n= 51).

|  | Placebo<br>n = 25 | Apremilast<br>n = 26 | P-value<sup>1</sup> |
| --- | --- | --- | --- |
| Sex (self-identified) |  |  |  |
| Male | 16 (64.0%) | 11 (42.3%) | 0.12 |
| Female | 9 (36.0%) | 15 (57.7%) |  |
| Race (self-identified) |  |  |  |
| American Indian/Alaska native | 0 (0%) | 1 (3.8%) | 0.27 |
| Asian | 0 (0%) | 0 (0%) |  |
| Black/African American | 0 (0%) | 2 (7.7%) |  |
| Native Hawaiian/other Pacific Islander | 1 (4.0%) | 0 (0%) |  |
| White | 23 (92.0%) | 20 (76.9%) |  |
| > 1 Race | 1 (4.0%) | 0 (0%) |  |
| Unknown/ Decline | 0 (0%) | 0 (0%) |  |
| Ethnicity (self-identified) |  |  |  |
| Hispanic/Latino | 2 (4.0%) | 5 (19.2%) | 0.24 |
| Not Hispanic/Latino | 24 (96.0%) | 20 (76.9%) |  |
| Age, years | 43.3 (18.5) | 39.10 (13.9) | 0.37 |
| BMI | 26.2 (4.0) | 26.3 (4.9) | 0.97 |
| C-Reactive Protein |  |  |  |
| Normal (<2mg/L) | 18 (72.0%) | 21 (80.8%) | 0.46 |
| High (≥2mg/L) | 7 (28.0%) | 5 (19.2%) |  |
| DSM V Symptoms, number<sup>2</sup> | 6.2 (2.4) | 6.6 (2.2) | 0.60 |
| Alcohol Use History |  |  |  |
| Age of 1<sup>st</sup> drink, years | 14.3 (2.9) | 14.5 (3.7) | 0.82 |
| Age of onset of alcohol-related problems, years | 23.5 (8.8) | 21.9 (6.7) | 0.47 |
| Years of heavy drinking<sup>3</sup> | 11.1 (10.1) | 13.4 (10.9) | 0.43 |
| Baseline Drinking (76d pre) |  |  |  |
| Drinks/Drinking Day | 5.2 (2.3) | 5.6 (2.8) | 0.64 |
| Drinks/Day<sup>4</sup> | 4.3 (1.6) | 4.4 (1.9) | 0.98 |
| % Drinking Days | 86 (0.2) | 84 (0.2) | 0.76 |
| % Heavy Drinking Days<sup>3</sup> | 45 (0.2) | 46 (0.3) | 0.95 |

<sup>1</sup> All values are means (standard deviations) unless otherwise specified.
<sup>2</sup> Numbers >5 indicate severe AUD.
<sup>3</sup> Heavy drinking is defined as ≥5 drinks/d (males), ≥4 drinks/d (females).
<sup>4</sup> Includes non-drinking days in mean (sd).

**Extended Data Table 2.** Pre- and post-treatment physiological indicators of treatment response.

| Characteristic | Placebo<br>n= 25 | n | Apremilast<br>n= 26 | n | P-value |
| --- | --- | --- | --- | --- | --- |
| **Apremilast in plasma (ng/mL)** | 0.0 (0.0) | 25 | 131.0 (210.8) | 21 | n/a |
| **Cytokines (pg/mL)** |  |  |  |  |  |
| **TNF-α** |  |  |  |  |  |
| Pre | 6.4 (1.9) | 20 | 6.0 (1.9) | 20 | 0.42 |
| Post | 6.4 (1.9) |  | 6.0 (1.5) |  | 0.24 |
| Change | 0.09 (1.2) |  | 0.10 (1.2) |  | 0.97 |
| **CCL2** |  |  |  |  |  |
| Pre | 320.8 (77.8) | 20 | 281.6 (98.5) | 20 | 0.17 |
| Post | 303.0 (77.1) |  | 268.1 (87.0) |  | 0.18 |
| Change | -13.6 (62.7) |  | -18.5 (67.4) |  | 0.81 |
| **CXCL10** |  |  |  |  |  |
| Pre | 51.3 (25.7) | 20 | 53.7 (40.7) | 20 | 0.82 |
| Post | 51.9 (24.5) |  | 54.1 (41.0) |  | 0.84 |
| Change | -0.3 (10.1) |  | -5.8 (17.1) |  | 0.23 |
| **HPA-axis (pg/mL)** |  |  |  |  |  |
| **Cortisol** |  |  |  |  |  |
| Pre | 711.9 (560.7) | 21 | 946.5 (861.5) | 22 | 0.29 |
| Post | 504.6 (241.2) |  | 866.0 (834.3) |  | 0.06 |
| Change | -207.3 (496.3) |  | -80.5 (856.0) |  | 0.72 |
| **Endotoxin (EU/ml)** |  |  |  |  |  |
| **LPS** |  |  |  |  |  |
| Pre | 0.10 (0.19) | 12 | 0.07 (0.15) | 11 | 0.62 |
| Post | 0.13 (0.25) |  | 0.06 (0.15) |  | 0.48 |
| Change | 0.02 (0.32) |  | 0.00 (0.02) |  | 0.78 |
| **Liver Function Tests (IU/L)** |  |  |  |  |  |
| **GGT** |  |  |  |  |  |
| Pre | 30.5 (29.1) | 21 | 25.0 (17.2) | 21 | 0.46 |
| Post | 29.3 (32.0) |  | 23.1 (14.8) |  | 0.43 |
| Change | -1.19 (8.6) |  | -2.24 (6.6) |  | 0.66 |
| **AST** |  |  |  |  |  |
| Pre | 23.8 (8.5) | 21 | 24.2 (9.8) | 21 | 0.87 |
| Post | 23.5 (10.3) |  | 20.9 (7.11) |  | 0.34 |
| Change | -0.24 (9.0) |  | -3.71 (9.83) |  | 0.24 |
| **ALT** |  |  |  |  |  |
| Pre | 22.9 (13.9) | 21 | 22.5 (12.8) | 21 | 0.91 |
| Post | 25.3 (18.2) |  | 20.8 (12.2) |  | 0.34 |
| Change | 2.43 (8.5) |  | -2.3 (9.8) |  | 0.10 |

Notes: Random missing data reflects a blood draw problem due to dehydration, or a value occurring below the limit of detectability. Determination of serum LPS was initiated in the latter half (n= 23) of the study.

**Extended Data Table 3.** Adverse events occurring in ≥ 5% of subjects.

| Body system | Event | Placebo<br>n = 25 | Apremilast<br>n = 26 |
| --- | --- | --- | --- |
| Gastrointestinal | Abdominal pain | 0 (0%) | 4 (15.4%) |
|  | Diarrhea | 4 (16.0%) | 9 (34.6%) |
|  | Dyspepsia | 1 (4.0%) | 1 (3.8%) |
|  | Nausea | 4 (16.0%) | 11 (42.3%) |
| General disorders | Hangover | 3 (12.0%) | 4 (15.4%) |
| Musculoskeletal | Back pain | 3 (12.0%) | 2 (7.7%) |
|  | Myalgia | 1 (4.0%) | 1 (3.8%) |
| Nervous system | Headache | 9 (36.0%) | 10 (38.5%) |
|  | Somnolence | 3 (12.0%) | 7 (26.9%) |
| Psychiatric | Insomnia | 2 (8.0%) | 2 (7.7%) |
| Respiratory | Allergic rhinitis | 1 (4.0%) | 1 (3.8%) |

**Extended data Table 4:** **Membrane properties and action potential characteristics of NAc medium spiny neurons (MSNs) treated with apremilast (1 µM) or vehicle (0.002% DMSO).** There were no statistically significant Treatment or Treatment x Cell Type interaction effects for any measures in this table. &: p < 0.05, main effect of Cell Type by 2-way ANOVA. N: number of neurons; sample sizes vary between measures because not all neurons fired action potentials. E_m_: membrane potential; AP: action potential, fAHP: fast afterhyperpolarization; mAHP: medium AHP; sAHP: slow AHP.

|  | Treatment | MSN Type | n | Mean | SEM |  |
| --- | --- | --- | --- | --- | --- | --- |
| Resting $E_m$ (mV) | Vehicle | D1 | 19 | -71.19 | 0.71 | |
|  |  | D2 | 23 | -70.99 | 0.77 |  |
|  | Apremilast | D1 | 24 | -71.92 | 0.72 |  |
|  |  | D2 | 21 | -72.71 | 0.65 |  |
| $R_{in}$ (MΩ) | Vehicle | D1 | 19 | 122.0 | 12.4 | |
|  |  | D2 | 23 | 126.2 | 8.0 |  |
|  | Apremilast | D1 | 24 | 117.8 | 9.2 |  |
|  |  | D2 | 21 | 134.0 | 12.1 |  |
| Rheobase (pA) | Vehicle | D1 | 19 | 257.9 | 19.2 | & |
|  |  | D2 | 23 | 217.4 | 15.6 |  |
|  | Apremilast | D1 | 24 | 239.6 | 18.5 |  |
|  |  | D2 | 20 | 205.0 | 21.1 |  |
| AP Amplitude (mV) | Vehicle | D1 | 19 | 48.95 | 2.65 |  |
|  |  | D2 | 23 | 48.64 | 2.00 |  |
|  | Apremilast | D1 | 24 | 53.05 | 1.57 |  |
|  |  | D2 | 20 | 48.15 | 1.58 |  |
| AP Half-Width (ms) | Vehicle | D1 | 19 | 1.65 | 0.08 |  |
|  |  | D2 | 23 | 1.59 | 0.05 |  |
|  | Apremilast | D1 | 24 | 1.58 | 0.05 |  |
|  |  | D2 | 20 | 1.64 | 0.05 |  |
| fAHP Amplitude (mV) | Vehicle | D1 | 19 | -9.81 | 1.30 |  |
|  |  | D2 | 23 | -10.39 | 1.01 |  |
|  | Apremilast | D1 | 24 | -9.87 | 1.00 |  |
|  |  | D2 | 20 | -8.70 | 0.66 |  |
| mAHP Amplitude (mV) | Vehicle | D1 | 19 | -9.56 | 0.63 | & |
|  |  | D2 | 23 | -7.79 | 0.43 |  |
|  | Apremilast | D1 | 24 | -8.64 | 0.42 |  |
|  |  | D2 | 20 | -8.17 | 0.48 |  |
| sAHP Amplitude (mV) | Vehicle | D1 | 19 | -8.51 | 0.57 | & |
|  |  | D2 | 23 | -6.29 | 0.53 |  |
|  | Apremilast | D1 | 24 | -7.44 | 0.47 |  |
|  |  | D2 | 20 | -7.24 | 0.54 |  |
| Max AP frequency (Hz) | Vehicle | D1 | 16 | 45.74 | 3.53 |  |
|  |  | D2 | 20 | 46.10 | 3.69 |  |
|  | Apremilast | D1 | 20 | 39.47 | 2.79 |  |
|  |  | D2 | 18 | 40.27 | 3.36 |  |

**Extended data Figure 1:**
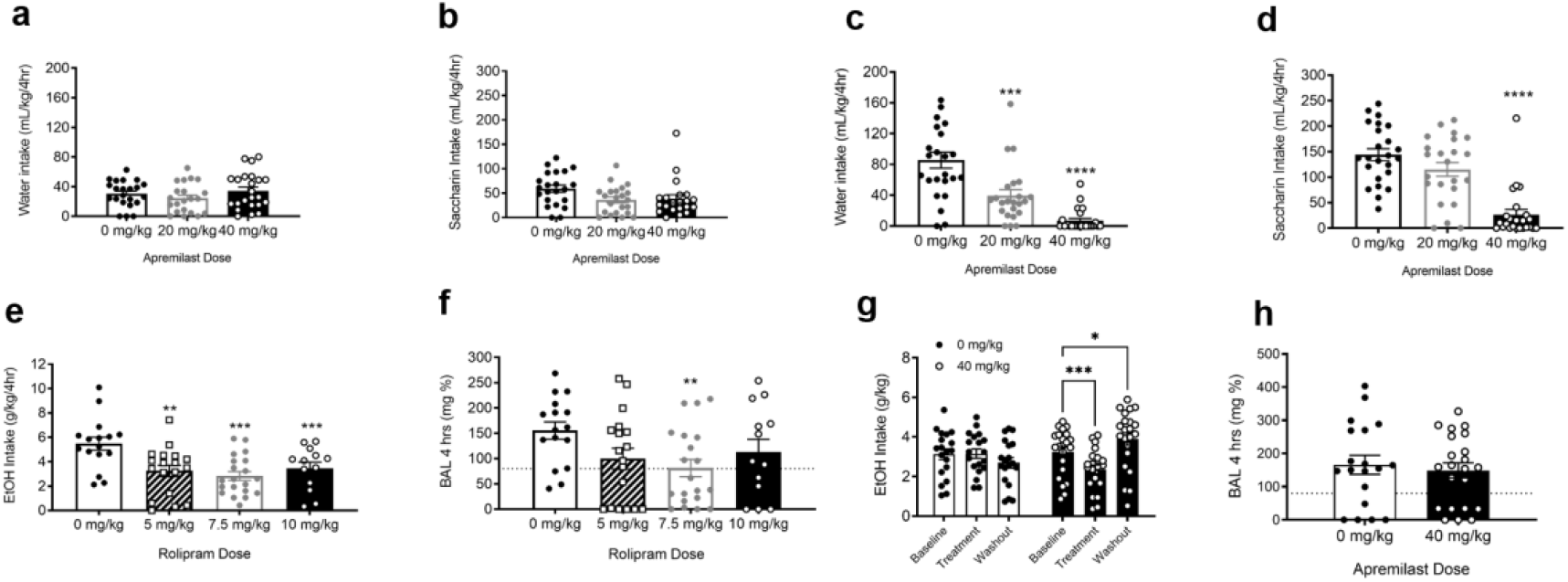
Acute and chronic PDE4 inhibition reduces binge-like ethanol drinking in genetic risk models. **A**, Water intake (mL/kg/4hrs) for HDID-1 (n = 10-12/sex/apremilast treatment); no main effects [F(2,63) = 1.27; p > 0.05]. **b**, Saccharin intake (mL/kg/4hrs) for HDID-1; no main effects [F(2,62) = 3.10; p > 0.05]. **c**, Water intake (mL/kg/4hrs) for HDID-2 (n = 11-12/sex/apremilast treatment); main effect of treatment [F(2,68) = 26.77; p < 0.0001], 20 and 40 mg/kg reduced water intake compared to 0 mg/kg. **d**, Saccharin intake (mL/kg/4hrs) for HDID-2; main effect of treatment [F(2,68) = 27.82; p < 0.0001], 40 mg/kg apremilast reduced saccharin intake compared to 0 mg/kg. **e**, Binge-like ethanol intake (g/kg/4hrs) for HDID-1 (n = 13-20/rolipram treatment); main effect of rolipram [F(3,84) = 10.40; p < 0.001], with no main effects of sex or sex X treatment interaction; all three doses of rolipram reduced ethanol intake in HDID-1 mice. **F**, Blood alcohol levels (mg %); main effect rolipram [F(3,84) = 10.40; p < 0.001], with no main effects of sex or sex X treatment interaction; 7.5 mg/kg of rolipram significantly reduced BALs. **G**, Average 4-hr ethanol intake over 6-week test (Wk 1: Baseline; Wks 2-5: treatment; Wk 6: Washout) for apremilast treated HDID-1 mice (n = 10-12/sex/apremilast treatment); main effect of time [F(2,78) = 5.68; p < 0.01] and a time by treatment interaction [F(2,78) = 17.56; p < 0.0001]; 40 mg/kg reduced ethanol intake compared to baseline and washout intake was higher than baseline. **H**, Blood alcohol levels (mg %) for end of week-6, 4-hr drinking; no main effects. (* = *p* < 0.05, ** = p < 0.005, *** = *p* < 0.001, **** = *p* < 0.0001). Dashed line indicates level of intoxication (80 mg %).

**Extended Data Figure 2:**
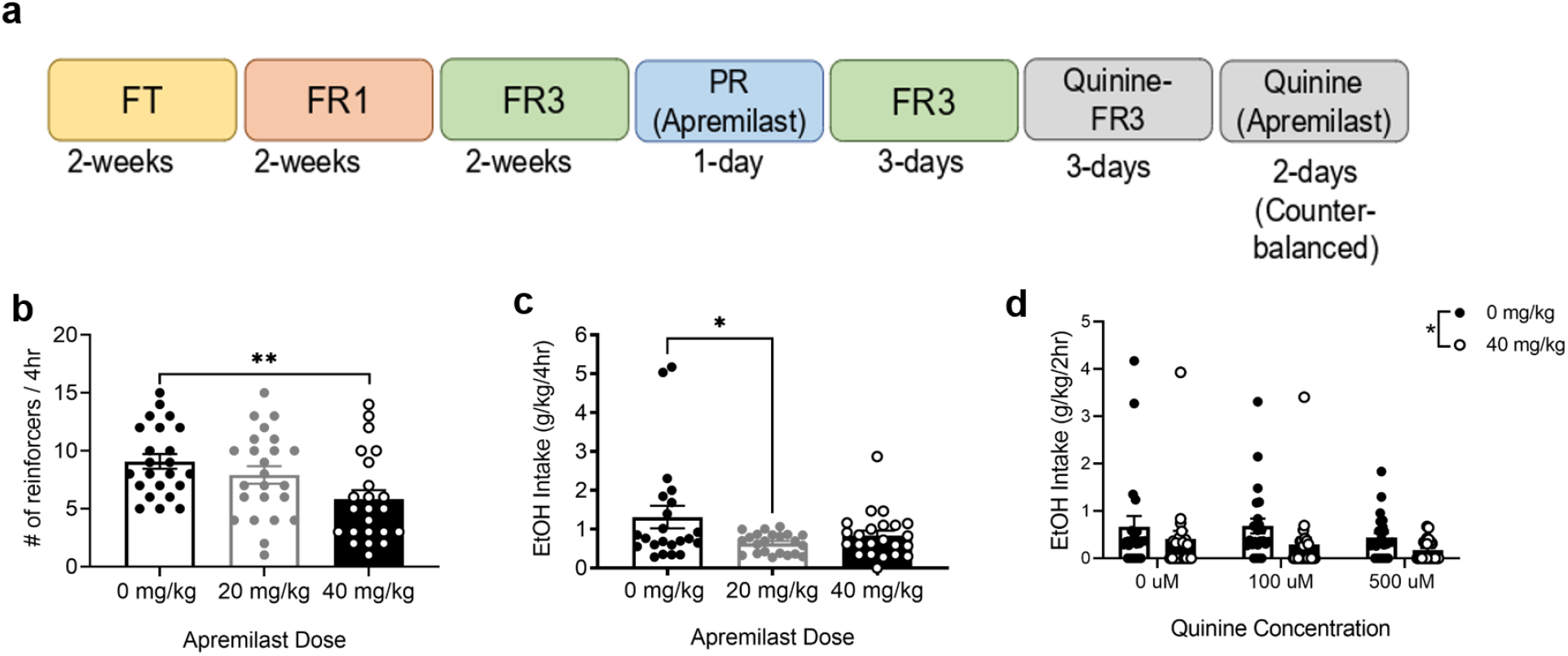
Apremilast reduces the motivation for ethanol in iHDID-1 mice. **A**, Experimental timeline of operant ethanol self-administration testing [Food training (FT); Fixed Ratio-1 and -3 (FR1, FR3); Progressive Ratio (PR)]. **B**, Average number of ethanol reinforcers during 4-hr for iHDID-1 mice (n =10-12/sex/apremilast treatment), PR session; main effect of apremilast treatment [F(2,64) = 6.63; p < 0.01] and sex [F(1,64) = 13.0; p < 0.001], with no treatment X sex interaction; 40 mg/kg apremilast reduced the reinforcers earned in iHDID-1 mice. **C**, Ethanol intake (g/kg/4hrs) during PR; main effect of apremilast treatment [F(2,61) = 3.87; p < 0.05], with no effect of sex or treatment X sex interaction. 20 mg/kg of apremilast reduced intake in iHDID-1 mice. **D**, Ethanol intake (g/kg/2hrs) during quinine–adulterated FR3 sessions; main effect of apremilast treatment [F(1, 64) = 6.97; p < 0.05] and a treatment X sex X quinine interaction [F(1,64) = 5.51; p < 0.01], with no effect of sex or a treatment x sex interaction; 40 mg/kg apremilast reduced quinine-adulterated ethanol intake. (* = *p* < 0.05, ** = *p* < 0.005).

**Extended Data Figure 3:**
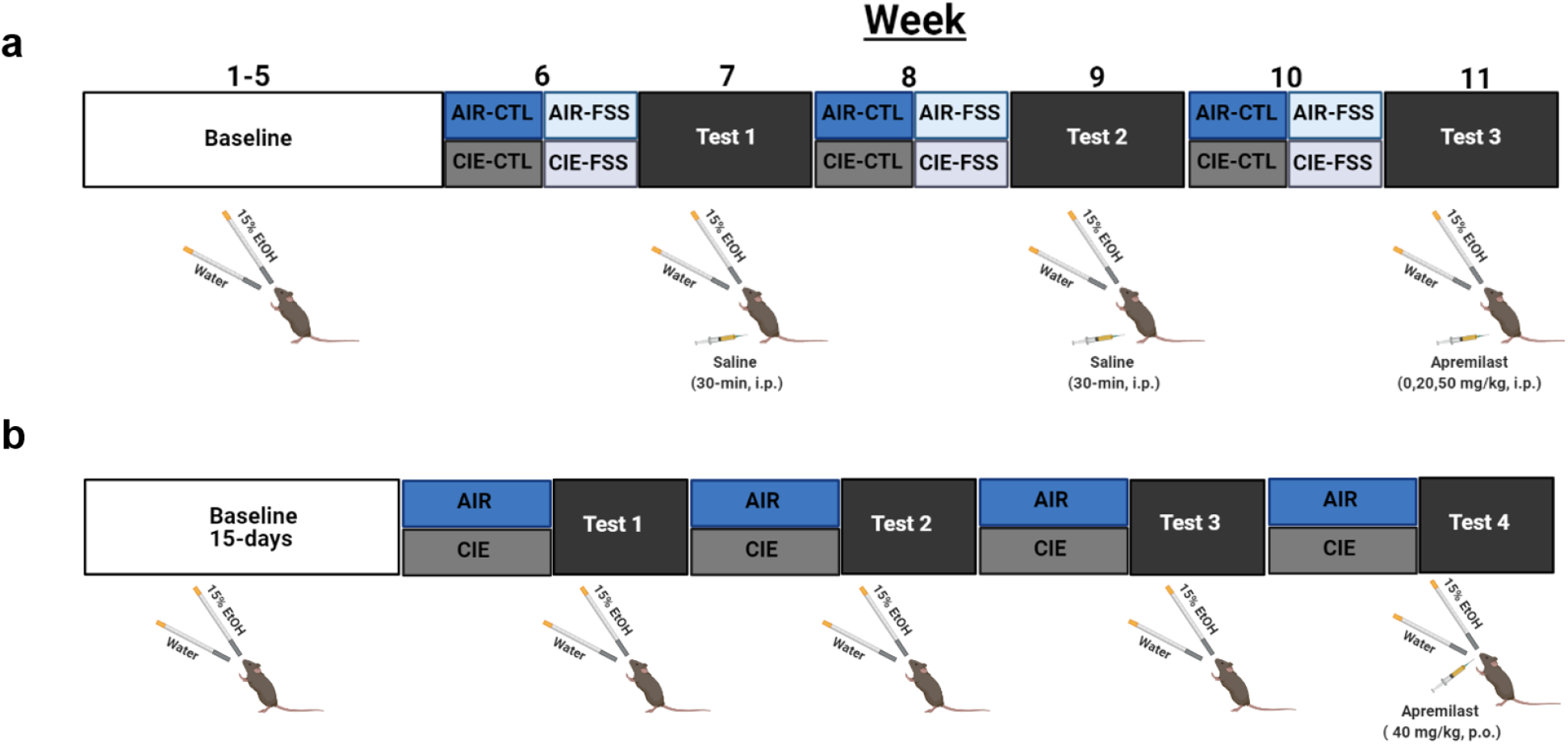
Experimental timelines for dependence induced escalations in binge-like drinking in C57BL/6J mice. **A**, Stress-CIE: Female and male C57BL/6J mice were given limited access to 15% ethanol and water for 2-hr, beginning 30-min prior to start of the dark cycle for 5-weeks (5 days/week; Baseline). Mice then received air or vapor exposure for 16 hrs/day for 4-days for 3-cycles (weeks 6,8, and 10). Mice were further divided into control (CTL) and forced swim stress (FSS) groups, whereby mice experienced FSS (10-min) 4-hr prior to DID during tests 1-3 (weeks 7, 9, and 11). Mice received apremilast (0,20,40 mg/kg), 30-min prior to drinking on test 3. **B**, CIE: Female and male C57BL/6J mice were given limited access to water and 15% ethanol for 2-hr, beginning 30-min prior to start of the dark cycle for 15 days (5 days/week; Baseline). Mice were then exposed to air or ethanol vapor for 16 hrs/day for 4 followed by limited access to water and 15% ethanol (72-hrs later) for 4 cycles. Mice then received an oral gavage of apremilast (40 mg/kg) 2-hrs prior to drinking access.

**Extended Data Figure 4.**
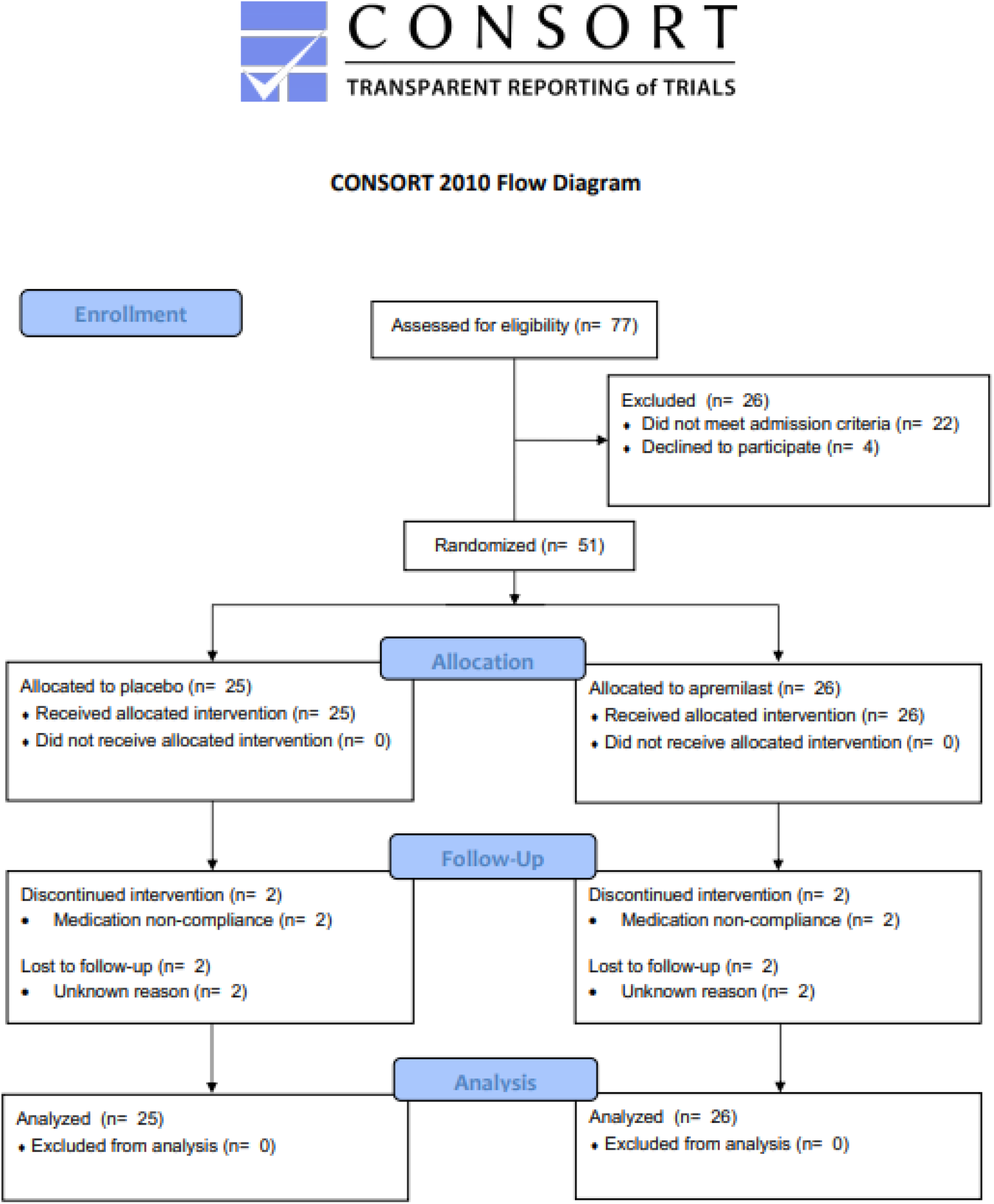
CONSORT 2010 flow diagram of phase IIa double-blind, placebo-controlled proof-of-concept (POC) study.

**Extended Data Figure 5:**
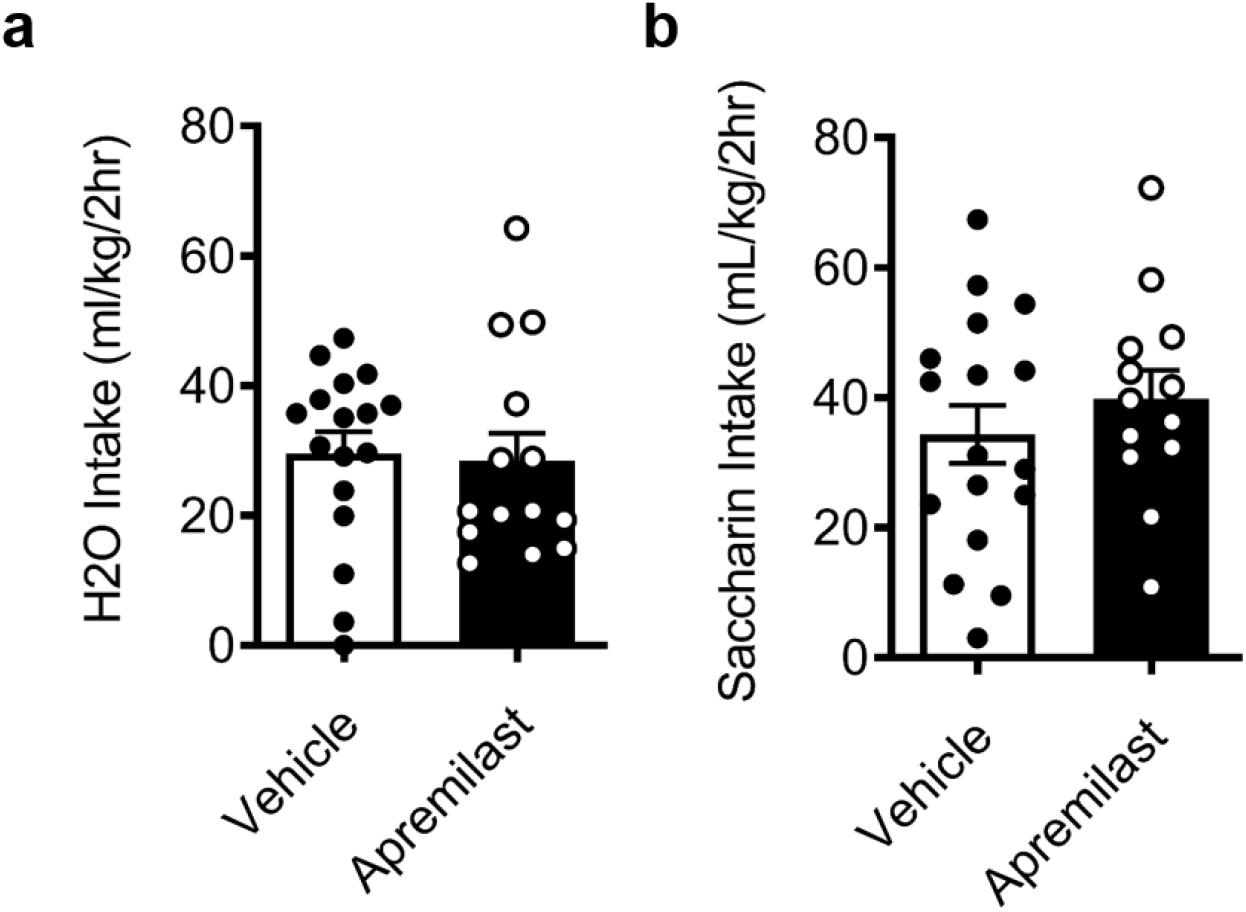
Intra-NAc Apremilast has no effect on water or saccharin intake in HDID-1 mice. **A**, Water intake (mL/kg/2hrs) following intra-NAc apremilast infusions (0 or 2 µg/µL/side), no main effect (Student’s t-test; p > 0.05). **b**, Saccharin intake (mL/kg/2hrs) following intra-NAc apremilast infusions (0 or 2 µg/µL/side), no main effect (Student’s t-test; p > 0.05).

## References cited

1. Prevention, C. for D. C. and. Excessive drinking is draining the US economy. Centers Dis. Control Prev. (2016).

2. Egli, M. Advancing pharmacotherapy development from preclinical animal studies. in The Neuropharmacology of Alcohol 537–578 (Springer, 2018).

3. Akbar, M., Egli, M., Cho, Y. E., Song, B. J. & Noronha, A. Medications for alcohol use disorders: An overview. Pharmacology and Therapeutics (2018) doi:10.1016/j.pharmthera.2017.11.007.

4. Robinson, G. et al. Neuroimmune pathways in alcohol consumption: Evidence from behavioral and genetic studies in rodents and humans. in International Review of Neurobiology (2014). doi:10.1016/B978-0-12-801284-0.00002-6.

5. Crews, F. T., Lawrimore, C. J., Walter, T. J. & Coleman, L. G. The role of neuroimmune signaling in alcoholism. Neuropharmacology (2017) doi:10.1016/j.neuropharm.2017.01.031.

6. Varodayan, F. P. et al. Role of TLR4 in the modulation of central amygdala GABA transmission by CRF following restraint stress. Alcohol Alcohol. 53, 642–649 (2018).

7. Clarke, T. K. et al. Genome-wide association study of alcohol consumption and genetic overlap with other health-related traits in UK biobank (N=112117). Mol. Psychiatry (2017) doi:10.1038/mp.2017.153.

8. Liu, M. et al. Association studies of up to 1.2 million individuals yield new insights into the genetic etiology of tobacco and alcohol use. Nature Genetics (2019) doi:10.1038/s41588-018-0307-5.

9. Seyhan, A. A. Lost in translation: the valley of death across preclinical and clinical divide – identification of problems and overcoming obstacles. Transl. Med. Commun. (2019) doi:10.1186/s41231-019-0050-7.

10. Nadeau, J. H. & Auwerx, J. The virtuous cycle of human genetics and mouse models in drug discovery. Nat. Rev. Drug Discov. 18, 255–272 (2019).

11. Rhodes, J. S., Best, K., Belknap, J. K., Finn, D. A. & Crabbe, J. C. Evaluation of a simple model of ethanol drinking to intoxication in C57BL/6J mice. Physiol. Behav. (2005) doi:10.1016/j.physbeh.2004.10.007.

12. Jensen, B. E. et al. Ethanol-Related Behaviors in Mouse Lines Selectively Bred for Drinking to Intoxication. Brain Sci. 11, 189 (2021).

13. Sneddon, E. A., Ramsey, O. R., Thomas, A. & Radke, A. K. Increased responding for alcohol and resistance to aversion in female mice. Alcohol. Clin. Exp. Res. 44, 1400– 1409 (2020).

14. Seif, T. et al. Cortical activation of accumbens hyperpolarization-active NMDARs mediates aversion-resistant alcohol intake. Nat. Neurosci. 16, 1094–1100 (2013).

15. Hopf, F. W. & Lesscher, H. M. B. Rodent models for compulsive alcohol intake. Alcohol 48, 253–264 (2014).

16. Anderson, R. I., Lopez, M. F. & Becker, H. C. Stress-induced enhancement of ethanol intake in C57Bl/6J mice with a history of chronic ethanol exposure: Involvement of kappa opioid receptors. Front. Cell. Neurosci. (2016) doi:10.3389/fncel.2016.00045.

17. Lopez, M. F., Anderson, R. I. & Becker, H. C. Effect of different stressors on voluntary ethanol intake in ethanol-dependent and nondependent C57BL/6J mice. Alcohol 51, 17– 23 (2016).

18. Jun Xu, Hsiao-Yen Ma, Xiao Liu, Sara Rosenthal, Jacopo Baglieri, Ryan McCubbin, Mengxi Sun, Yukinori Koyama, Cedric G. Geoffroy, Kaoru Saijo, Linshan Shang, Takahiro Nishio, Igor Maricic, Max Kreifeldt, Praveen Kusumanchi, Amanda Roberts, Binhai Zheng, V., and T. K. Blockade of IL-17 signaling reverses alcohol-induced liver injury and excessive alcohol drinking in mice. JCI insight 5, (2020).

19. Koob, G. F. & Volkow, N. D. Neurocircuitry of addiction. Neuropsychopharmacology (2010) doi:10.1038/npp.2009.110.

20. Koob, G. F. & Volkow, N. D. Neurobiology of addiction: a neurocircuitry analysis. The Lancet Psychiatry (2016) doi:10.1016/S2215-0366(16)00104-8.

21. Henderson, M. B. et al. Deep brain stimulation of the nucleus accumbens reduces alcohol intake in alcohol-preferring rats. Neurosurg. Focus 29, E12 (2010).

22. Heinze, H. J. et al. Counteracting incentive sensitization in severe alcohol dependence using deep brain stimulation of the nucleus accumbens: Clinical and basic science aspects. Front. Hum. Neurosci. (2009) doi:10.3389/neuro.09.022.2009.

23. Purohit, K. et al. Pharmacogenetic Manipulation of the Nucleus Accumbens Alters Binge-Like Alcohol Drinking in Mice. Alcohol. Clin. Exp. Res. (2018) doi:10.1111/acer.13626.

24. Townsley, K. G., Borrego, M. B. & Ozburn, A. R. Effects of chemogenetic manipulation of the nucleus accumbens core in male C57BL/6J mice. Alcohol 91, 21–27 (2021).

25. Crabbe, J. C. Using signatures of directional selection to guide discovery. in Molecular-Genetic and Statistical Techniques for Behavioral and Neural Research (2018). doi:10.1016/B978-0-12-804078-2.00011-8.

26. Iancu, O. D. et al. Selection for drinking in the dark alters brain gene coexpression networks. Alcohol. Clin. Exp. Res. (2013) doi:10.1111/acer.12100.

27. Ferguson, L. B. et al. Genome-Wide Expression Profiles Drive Discovery of Novel Compounds that Reduce Binge Drinking in Mice. Neuropsychopharmacology (2018) doi:10.1038/npp.2017.301.

28. Blednov, Y. A., Benavidez, J. M., Black, M. & Adron Harris, R. Inhibition of phosphodiesterase 4 reduces ethanol intake and preference in C57BL/6J mice. Front. Neurosci. (2014) doi:10.3389/fnins.2014.00129.

29. Blednov, Y. A., Da Costa, A. J., Harris, R. A. & Messing, R. O. Apremilast alters behavioral responses to ethanol in mice: II. Increased sedation, intoxication, and reduced acute functional tolerance. Alcohol. Clin. Exp. Res. 42, 939–951 (2018).

30. Li, Z. et al. Intermittent high-dose ethanol exposures increase motivation for operant ethanol self-administration: possible neurochemical mechanism. Brain Res. 1310, 142– 153 (2010).

31. Weiss, F., Lorang, M. T., Bloom, F. E. & Koob, G. F. Oral alcohol self-administration stimulates dopamine release in the rat nucleus accumbens: Genetic and motivational determinants. J. Pharmacol. Exp. Ther. (1993).

32. Becker, H. C. & Lopez, M. F. Increased ethanol drinking after repeated chronic ethanol exposure and withdrawal experience in C57BL/6 mice. Alcohol. Clin. Exp. Res. 28, 1829– 1838 (2004).

33. Finn, D. A. et al. Increased drinking during withdrawal from intermittent ethanol exposure is blocked by the CRF receptor antagonist D-Phe-CRF(12-41). Alcohol. Clin. Exp. Res. (2007) doi:10.1111/j.1530-0277.2007.00379.x.

34. Gruol, D. L., Melkonian, C., Huitron-Resendiz, S. & Roberts, A. J. Alcohol alters IL-6 signal transduction in the CNS of transgenic mice with increased astrocyte expression of IL-6. Cell. Mol. Neurobiol. 1–18 (2020).

35. Gruol, D. L., Huitron-Resendiz, S. & Roberts, A. J. Altered brain activity during withdrawal from chronic alcohol is associated with changes in IL-6 signal transduction and GABAergic mechanisms in transgenic mice with increased astrocyte expression of IL-6. Neuropharmacology 138, 32–46 (2018).

36. Papp, K. et al. Apremilast, an oral phosphodiesterase 4 (PDE4) inhibitor, in patients with moderate to severe plaque psoriasis: results of a phase III, randomized, controlled trial (Efficacy and Safety Trial Evaluating the Effects of Apremilast in Psoriasis [ESTEEM] 1). J. Am. Acad. Dermatol. 73, 37–49 (2015).

37. Schafer, P. H. et al. Apremilast, a cAMP phosphodiesterase-4 inhibitor, demonstrates anti-inflammatory activity in vitro and in a model of psoriasis. Br. J. Pharmacol. 159, 842–855 (2010).

38. Association, A. P. Diagnostic and statistical manual of mental disorders (DSM-5®). (American Psychiatric Pub, 2013).

39. Laird, N. M. & Ware, J. H. Random-effects models for longitudinal data. Biometrics 963– 974 (1982).

40. Bolker, B., Skaug, H., Magnusson, A. & Nielsen, A. Getting started with the glmmADMB package. (2012).

41. Cohen, J. Statistical power analysis for the behavioral sciences. (Academic press, 2013).

42. Maisel, N. C., Blodgett, J. C., Wilbourne, P. L., Humphreys, K. & Finney, J. W. Meta-analysis of naltrexone and acamprosate for treating alcohol use disorders: when are these medications most helpful? Addiction 108, 275–293 (2013).

43. Nishi, A. et al. Distinct roles of PDE4 and PDE10A in the regulation of cAMP/PKA signaling in the striatum. J. Neurosci. 28, 10460–10471 (2008).

44. Voges, J., Müller, U., Bogerts, B., Münte, T. & Heinze, H. J. Deep brain stimulation surgery for alcohol addiction. World Neurosurgery (2013) doi:10.1016/j.wneu.2012.07.011.

45. Kuhn, J. et al. Successful deep brain stimulation of the nucleus accumbens in severe alcohol dependence is associated with changed performance monitoring. Addict. Biol. (2011) doi:10.1111/j.1369-1600.2011.00337.x.

46. Pozhidayeva, D. Y. et al. Chronic chemogenetic stimulation of the nucleus accumbens produces lasting reductions in binge drinking and ameliorates alcohol-related morphological and transcriptional changes. Brain Sci. (2020) doi:10.3390/brainsci10020109.

47. Kircher, D. M., Aziz, H. C., Mangieri, R. A. & Morrisett, R. A. Ethanol experience enhances glutamatergic ventral hippocampal inputs to D1 receptor-expressing medium spiny neurons in the nucleus accumbens shell. J. Neurosci. 39, 2459–2469 (2019).

48. Renteria, R., Maier, E. Y., Buske, T. R. & Morrisett, R. A. Selective alterations of NMDAR function and plasticity in D1 and D2 medium spiny neurons in the nucleus accumbens shell following chronic intermittent ethanol exposure. Neuropharmacology 112, 164–171 (2017).

49. Bell, R. L. et al. Ibudilast reduces alcohol drinking in multiple animal models of alcohol dependence. Addict. Biol. 20, 38–42 (2015).

50. Ray, L. A. et al. Development of the neuroimmune modulator ibudilast for the treatment of alcoholism: a randomized, placebo-controlled, human laboratory trial. Neuropsychopharmacology 42, 1776–1788 (2017).

51. Burnette, E. M., Baskerville, W.-A., Grodin, E. N. & Ray, L. A. Ibudilast for alcohol use disorder: study protocol for a phase II randomized clinical trial. Trials 21, 1–19 (2020).

52. Hu, W. et al. Inhibition of phosphodiesterase-4 decreases ethanol intake in mice. Psychopharmacology (Berl). (2011) doi:10.1007/s00213-011-2290-8.

53. Blednov, Y. A. et al. Apremilast Alters Behavioral Responses to Ethanol in Mice: I. Reduced Consumption and Preference. Alcohol. Clin. Exp. Res. (2018) doi:10.1111/acer.13616.

54. Liu, X. et al. The phosphodiesterase-4 inhibitor roflumilast decreases ethanol consumption in C57BL/6J mice. Psychopharmacology (Berl). 234, 2409–2419 (2017).

55. Ozburn, A. R. et al. Effects of Pharmacologically Targeting Neuroimmune Pathways on Alcohol Drinking in Mice Selectively Bred to Drink to Intoxication. Alcohol. Clin. Exp. Res. (2020) doi:10.1111/acer.14269.

56. Wen, R. T. et al. The Phosphodiesterase-4 (PDE4) Inhibitor Rolipram Decreases Ethanol Seeking and Consumption in Alcohol-Preferring Fawn-Hooded Rats. Alcohol. Clin. Exp. Res. (2012) doi:10.1111/j.1530-0277.2012.01845.x.

57. Farris, S. P. et al. Transcriptome Analysis of Alcohol Drinking in Non-Dependent and Dependent Mice Following Repeated Cycles of Forced Swim Stress Exposure. Brain Sci. 10, 275 (2020).

58. Koob, G. F. Neurocircuitry of alcohol addiction: Synthesis from animal models. In Handbook of Clinical Neurology (2014). doi:10.1016/B978-0-444-62619-6.00003-3.

59. Purohit, K. et al. Pharmacogenetic Manipulation of the Nucleus Accumbens Alters Binge-Like Alcohol Drinking in Mice. Alcohol. Clin. Exp. Res. (2018) doi:10.1111/acer.13626.

60. Strong, C. E. et al. Chemogenetic selective manipulation of nucleus accumbens medium spiny neurons bidirectionally controls alcohol intake in male and female rats. Sci. Rep. 10, 1–15 (2020).

61. Knapp, C. M., Tozier, L., Pak, A., Ciraulo, D. A. & Kornetsky, C. Deep brain stimulation of the nucleus accumbens reduces ethanol consumption in rats. Pharmacol. Biochem. Behav. 92, 474–479 (2009).

62. Blokland, A. et al. Phosphodiesterase Type 4 Inhibition in CNS Diseases. Trends Pharmacol. Sci. 40, 971–985 (2019).

