## Supplementary material for "The FDA-approved drug apremilast suppresses alcohol intake: clinical and pre-clinical validation": Materials and Methods

### Mouse strains and animal care

To test whether apremilast reduces binge-like drinking and the motivation for alcohol, we used selectively bred High Drinking in the Dark (HDID-1, HDID-2) and inbred -HDID-1 (iHDID) mice of both sexes. Female and male HDID-1 were also used to determine whether PDE4 inhibition in the NAc alone is sufficient to reduce binge-like ethanol drinking. The above lines were bred and maintained in the Portland VA Medical Center Veterinary Medical Unit[1,2].

To determine whether apremilast reduces drinking in models of ethanol dependence-induced escalation in drinking we tested adult male and female C57BL/6J mice in two important paradigms, Chronic Intermittent Ethanol-Forced Swim Stress (CIE-FSS) and Chronic Intermittent Ethanol (CIE)[3]. The CIE-FSS experiment was performed at the Charleston Alcohol Research Center at the Medical University of South Carolina. The more chronic CIE procedure was conducted at the Animal Models Core Facility at the La Jolla Alcohol Research center (Scripps Institute)[4,5]. For both experiments, subjects were maintained on a 12-hr modified light/dark cycle with ad libitum access to food and water throughout experimentation.

For electrophysiological experiments, B6.Cg-Tg(Drd1a-tdTomato)6Calak/J ("*Drd1a*-tdTomato", C57BL/6J congenic) mice[6] were bred and maintained at the University of Texas at Austin. The colony of *Drd1a*-tdTomato mice (initial breeding pairs obtained from The Jackson Laboratory, Stock No. 016204) was maintained by crosses in which only one parent (typically the male) was a carrier of the *Drd1a*-tdTomato transgene. Experimental mice were pair-housed in standard cages with Sani-Chips wood bedding (PJ Murphy, Forest Products, Montville, NJ, USA) and a cotton fiber nestlet (Ancare; Bellmore, NY, USA), in a temperature controlled room (~21°C) with a reverse 12hr light:12 hr dark cycle (lights off at 9:30 am). Mice had *ad libitum* access to standard chow (LabDiet® 5LL2 Prolab RMH 1800) and a single bottle of tap water, and cages were changed once weekly.

All procedures were approved by the local Institutional Animal Care and Use Committee and were conducted in accordance with the NIH Guidelines for the Care and Use of Laboratory Animals for that institution.

**Drugs**

For the Drinking in the Dark tests, mice were offered 20% ethanol (v/v, in tap water; 200 proof, DeCon Labs, King of Prussia, PA) and for the chronic intermittent ethanol (CIE) studies, mice were offered 15% ethanol (v/v, prepared with Ethyl alcohol 200 proof. Pharmco-AAPER, Brookfield, CT). For binge-like drinking studies with serial testing, mice were offered 9.2 mM saccharin sodium hydrate (in tap water; Sigma-Aldrich). For binge-like drinking studies, apremilast (Toronto Research Chemicals) and rolipram (Sigma-Aldrich) were suspended in Tween-80 (1.875% v/v) and saline and administered intraperitoneally (i.p.) in a volume of 10 mL/kg mouse body weight. For the chronic intermittent ethanol (CIE) study, apremilast was prepared in an identical manner and administered 10 mL/kg by oral gavage, 2 hours prior to measured drinking.  For the intra-NAc apremilast study, 1 µL of 2 µg/µL apremilast (dissolved in 35% DMSO and Dulbecco's phosphate buffered saline, DPBS) was administered bilaterally into the nucleus accumbens (where cannula placement was verified using methylene blue[7]). For electrophysiology, frozen aliquots of 50 mM apremilast in 100% dimethyl sulfoxide (DMSO; Fisher) were defrosted and added to artificial cerebral spinal fluid (ACSF), yielding final concentrations of 1 µM apremilast and 0.002% DMSO. Vehicle solutions for each study were prepared identically without the addition of rolipram or apremilast.

**Testing the effects of PDE4 inhibition on binge-like ethanol drinking**

*Drinking in the Dark (DID) Test:* As in Rhodes et al. (2005), mice were habituated to individual housing and a new sipper bottle for one week prior to testing[8]. For the ethanol DID assay, mice were offered a single bottle of 20% ethanol (v/v, in tap water), 3 hours after lights off. The general approach used was to measure fluid intake daily for either 2 hrs (first three days) or 4 hrs (fourth day); see specific experimental details below. Water was then restored for the remainder of the day (unmeasured). Details for each study follow below. Mice were pseudorandomly assigned to treatment groups based on their alcohol intake during the first 3 days of DID to achieve comparable drinking levels prior to testing on the fourth day. Treatment group assignment was fixed and did not change.

*Rolipram:* HDID-1 male and female (n = 13-20/rolipram treatment) mice were habituated to individual housing for one week as described above. Mice were then subjected to ethanol DID, where they received access to 20% ethanol for 2-hrs on days 1, 2, and 3. On day 4, mice were treated with 0, 5, 7.5, or 10 mg/kg rolipram 30-min prior to a 4-hr DID session. Immediately after the drinking session, a 20 µL peri-orbital sinus blood sample was collected for determination of blood alcohol level (BAL) by gas chromatography using methods described previously (Crabbe and Ozburn et al., 2017).

*Apremilast:* HDID-1 male and female mice (n = 10-12/sex/apremilast treatment) were habituated to individual housing for one week as described above. Mice were then subjected to 3 weeks of DID testing (4 days/week), where mice were offered 20% ethanol in week 1, water in week 2, and 9.2 mM saccharin in week 3. Ethanol consumption is determined by many factors, including palatability; therefore, we tested the effects of apremilast on intake of ethanol, as well as water and a sweet tastant solution (saccharin). In week 1, mice received access to 20% ethanol for 2-hrs on days 1, 2, and 3. On day 4, mice were treated with 0, 20, or 40 mg/kg apremilast 30-min prior to a 4-hr ethanol DID session. Immediately after the ethanol drinking session, a 20 µL peri-orbital sinus blood sample was collected for determination of BAL by gas chromatography using methods described previously[9]. Water intake in week 2 and 9.2 mM saccharin intake in week 3 were measured identically to ethanol, whereby volumes were measured for 2-hrs on days 1, 2, and 3. On day 4, mice were treated with 0, 20, or 40 mg/kg apremilast 30-min prior to a 4-hr water DID session.

**Effects of apremilast on the motivation for ethanol**

To determine whether PDE4 inhibition reduces the motivation for ethanol and/or aversion resistant responding for ethanol, we tested the effects of apremilast on oral ethanol self-administration during a 1) progressive ratio (PR) schedule of reinforcement and for 2) quinine-adulterated ethanol. Mice were trained to acquire lever pressing for a food pellet reinforcer (food training) prior to measuring for responding and ethanol intake during fixed ratio responding (FR1, FR3), and PR for access to 20% ethanol or quinine-adulterated 20% ethanol. In total, 72 mice [n = 10-12/sex/apremilast treatment (0, 20, 40 mg/kg apremilast)] were used.

One week prior to testing, mice were habituated to individual housing and sipper bottles (matching those used for operant self-administration). 24 operant testing chambers (Med Associates, VT, USA) were used, which were housed in light and sound attenuating boxes. Chambers were operated using MedPC IV software (Med Associates, VT, USA). Each chamber was outfitted for lever responding for food (trough) or liquid (sipper) reinforcement, and contained a trough for pellet delivery, an ethanol extender (retracted between reinforcers), a lickometer, two levers with cue lights above, and one house light.

An experimental timeline is provided in extended data figure 2a. Food training (FT) was used to ensure mice learned to lever-press for access to a reinforcer prior to ethanol self-administration, as shown in Jensen *et al*. 2021[10]. Mice started food restriction one day prior to training and were maintained at 85% of their free feeding weight throughout FT. In brief, mice were placed for 1-hr daily in an operant conditioning chamber equipped with a chow pellet dispenser and two levers, during which the house light (signaling availability of a reinforcer) and the cue light above the active lever were illuminated continuously. Training was under an FR1 schedule, whereby a single lever response (FR1) of the active lever press (left) delivered 1 chow pellet as a reinforcer. To avoid overtraining, mice were food trained until they reached 3 sessions earning ≥ 25 chow pellet reinforcers / session. All mice met criteria by 7 sessions of FT.

*Operant ethanol self-administration behavior*: iHDID mice were trained in 10, 2-hr sessions of fixed ratio 1 (FR1; 5 days/week for 2 weeks) and 10, 2-hr sessions of FR3 (5 days/week for 2 weeks). Ethanol intake for each 2-hr session, number of active and inactive lever presses, and number of ethanol access periods (reinforcers) earned were recorded. Stability in responding was defined as < 20% variance within the mean reinforcers earned within the last 3 sessions of FR3. Mice not exhibiting greater than a 2:1 active: inactive lever pressing ratio (to ensure responses are specific to the reinforcer) or stable responding during FR3 did not continue to PR or quine-adulterated sessions.

To test the role of PDE4 in the motivation for ethanol, iHDID mice exhibiting stable FR3 responding were treated with apremilast (0, 20, or 40 mg/kg) 1 hr prior to a single 4-hr session on a PR schedule for 20% ethanol reinforcement. PR response requirements increased stepwise by [$a_n = 1/8(2n^2 + (-1)^n + 7)$ (the pattern being 1, 2, 3, 5, 7, 10, 13, 17, 21, 26 etc…)]. The last responding ratio reached in the 4 hr session was defined as the breakpoint. In addition to the breakpoint, ethanol intake, number of active and inactive lever presses, and number of ethanol access periods (reinforcers) earned were recorded.

To determine whether apremilast reduces aversion-resistant responding for ethanol, the above female and male iHDID mice were tested for quinine-adulterated ethanol responding on an FR3 schedule, similar to the design of Sneddon *et al.*[11]. After completion of the PR test, mice returned to 3, 2-hr sessions of FR3 responding for 20% ethanol, followed by 3, 1-hr sessions of

FR3 for 20% + quinine (0, 100, and 500 µM). Mice were pseudorandomized into treatment groups based on the average responses and intake during the last 3 sessions. Using a counterbalanced design, mice were tested for quinine-adulterated ethanol responding during 2, 1-hr sessions (on an FR3 schedule). Here, half of the group received apremilast (40 mg/kg) 1-hr prior to testing on Day 1 (and vehicle on day 2), and the other half received apremilast on day 2 (with vehicle on Day 1). The 40 mg/kg was chosen because preliminary results suggested it was more efficacious than 20 mg/kg at reducing intake and reinforcers earned. The number of ethanol access periods (reinforcers) earned, ethanol intake, and number of active and inactive lever presses were recorded.

**The effect of Apremilast on models of ethanol dependence**

*Stress + Chronic intermittent stress (CIE):* An experimental timeline is provided in extended data figure 3A. Adult male C57BL/6J were trained to drink ethanol daily (Mon-Fri) for 1-hr/day starting three hours after lights off. Ethanol (15% v/v) and water (2-bottle choice; 2-BC), were available during this 1-hr access period. After 5-weeks of baseline drinking mice were separated into four groups (CTL, FSS, CIE, CIE+FSS) and entered the CIE ± FSS phase of the study. That is, weekly cycles of CIE/Air exposure (as described above) were alternated with weekly test drinking cycles, with half the mice receiving 10-min FSS exposure 4-hrs prior to the drinking sessions (remaining mice were not disturbed)[3,12]. Therefore, the four groups evaluated in this study were CTL, CIE, FSS (air-exposed mice that experienced FSS before drinking), and CIE+FSS (mice that experienced both CIE and FSS). During test cycle 3, these four groups were further separated into groups based on the apremilast dose condition (0, 20, 40 mg/kg; N= 9-10/group). During Baseline and the first two test cycles, all mice received vehicle (saline) injections (IP) 30-min before ethanol access. During test cycle 3, mice were injected with apremilast (20 or 40 mg/kg) or vehicle 30-min before the drinking sessions. Solutions were prepared fresh daily mixing saline with apremilast. Mice received IP injections at a 10 ml/kg volume.

*CIE:* An experimental timeline is provided in Extended Data Figure 3 b. To test the effects of apremilast on a more chronic model of dependence induced escalations in ethanol drinking, female and male C57BL/6J mice (n = 10/sex/group) underwent a standard CIE protocol[13–15]. In brief, baseline drinking consisted of 2-hr access to 15% ethanol and water, 5-days/week for 3-weeks (15-days total). Mice were grouped based on equal ethanol and water consumption, into two balanced treatment groups: CIE or air (Control). Concentrations of ethanol vapor were adjusted based on the blood alcohol levels of the mice and target blood alcohol levels were 150-200 mg/dL. The CIE group was injected with 1.75 g/kg EtOH and 68.1 mg/kg pyrazole (alcohol dehydrogenase inhibitor, Sigma-Aldrich) and the Control group received pyrazole only and all mice were placed in the chambers for 16 hr. After the fourth day of exposure, mice were left undisturbed for 72-hr in their home cages, followed by an identical 2-BC for 5-days. The above CIE and limited access drinking sessions were repeated for a total of 4 test cycles. Tail blood (20 µl) was taken on the 3rd day of each cycle to analyze blood alcohol levels (Analox Instruments, Lunenburg, MA, USA).

Following the fourth cycle of vapor exposure, mice were acclimated to the gavage procedure by 7 daily injections of vehicle, given at 2:00PM. In a pilot study, the vehicle decreased drinking in non-acclimated mice if we gave it before the drinking session; therefore, we acclimated the mice to the gavage procedure to minimize negative associations between gavage and ethanol. Two-bottle choice sessions continued through this week with the weekend off. On the following Monday, 2-hr ethanol drinking was recorded for a baseline measure, on Tuesday all mice received oral vehicle 2-hr before two bottle choice, and on Wednesday all mice received oral apremilast 2-hr before the drinking session.

**Clinical Research Methods**

*Regulatory Approvals and Trial Registration:* The study protocol was approved by the Scripps Research Institutional Review Board (protocol #16-6821), was conducted under an Investigator-initiated IND (#135813), and is registered on ClinicalTrials.gov (NCT03175549).

*Setting:* This single-site study was conducted in the outpatient clinical research unit of the Laboratory of Clinical Neuropsychopharmacology, Pearson Center for Alcoholism and Addiction Research, Department of Molecular Medicine, The Scripps Research Institute, La Jolla, CA 92037.

*Subject Characteristics:* IRB-approved print, social media and internet advertisements were used to recruit research volunteers. Subject accrual occurred between 11/01/2017 and 04/01/2020. Subjects were paid $50 for completing an initial evaluation, $75 for completing the randomization visit and $150 for completing the final treatment visit.

Subjects were males or females, 18-65 years of age, who met Diagnostic and Statistical Manual for Mental Disorders – Fifth Edition[16] (DSM-5) criteria for Alcohol Use Disorder (AUD) ≥ moderate severity, and who were not seeking treatment and willing to take daily oral medication for research purposes.  Subjects were not pregnant, did not have a urine drug screen positive for substances of abuse other than alcohol, were not taking disallowed medications and did not have significant medical or psychiatric disorders that would increase potential risk or interfere with study outcomes, as determined by the study physician's review of medical history, vital signs, routine urine and blood tests, electrocardiogram, and physical examination. The Mini International Neuropsychiatric Interview[17] (M.I.N.I.) was used to establish the categorical diagnosis and severity of AUD and to rule out exclusionary psychiatric disorders. The Clinical

Institute Withdrawal Assessment for Alcohol-Revised[18] (CIWA-Ar) was used to assess severity of alcohol withdrawal symptoms; subjects were required to have a negative breathalyzer reading and a CIWA-Ar score < 9 at randomization, to eliminate acute alcohol or withdrawal effects.

*Outcome Measures*

*Efficacy:* The Time Line Follow Back Interview[19] (TLFB) was the primary measure used to access daily intake of standard drinks consumed over the 11-day period of *ad libitum* drinking. A standard drink contains ~14 grams of alcohol, such at one 12 oz beer is equivalent to one 5 oz glass of wine and 1.5 oz of distilled spirits. A daily drinking diary was used to confirm TLFB data.

*Safety:* Adverse drug experiences were recorded on standardized case report forms that depicted each side effect complaint in terms of its onset, duration, severity, relation to study medication and clinical action. Vital signs, routine urine and blood tests, EKG and physical exam were conducted pre and post treatment to verify that no clinically significant changes from baseline had occurred.

*Medication Conditions:* Apremilast was purchased from a retail specialty pharmacy, over-encapsulated by Austin Compounding Pharmacy in Austin, Texas, and matched with identical placebo capsules. Double-blind study drug was given in a standard 7-day titration to 90 mg/d apremilast or matched placebo, administered orally as two 30 mg capsules (60 mg) of apremilast or identical placebo in the AM, and one 30 mg capsule of apremilast or identical placebo in the PM, taken with or without food. Double-blind study drug was packaged in a blistercard with the subject's study ID number and the day and time of each dose indicated on the blistercard. Study drugs, packaging and dosing regimen were identical to preserve the double-blind. All participants, care providers, and those assessing outcomes were blinded to the identity of the assigned medication until after the trial was completed. Medication adherence was verified with returned blistercard and pill count. Apremilast plasma determinations were obtained on the last day of dosing, frozen (-80º C) and analyzed in batch after study completion to verify correct medication assignment per the randomization code and ingestion of active medication and were examined for an association with outcome on an exploratory basis, as apremilast has no established therapeutic plasma level. Determination of apremilast concentration in plasma was made by High Performance Liquid Chromatography (HPLC) in the laboratory of Dr. Esther Maier, Drug Dynamics Institute, TherapeUTex, University of Texas – Austin.

*Physiological Indicators:* Apremilast is a selective PDE4 and TNF-α inhibitor that acts on immune system targets. Hence, plasma for determination of cytokines (TNF-alpha, CCL2, CXXL10) and cortisol concentration was obtained, frozen (-80º C) and assayed in batch after study completion for retrospective evaluation as potential physiological moderators of treatment response. Levels of endotoxin in serum were measured as a marker of leakage of the intestinal barrier. Results ruled out such leakage as an explanation for cytokine levels observed, thereby also ruling out endotoxins as potentially confounding treatment response to apremilast in AUD.

The following assays were run by the CSAR Cell and Immunology Core, P30 Center on Intersystem Regulation by Drugs of Abuse, DA013429. Katz School of Medicine at Temple University:

*Luminex Assay:* Plasma levels of chemokines and cytokines were determined using the Human Magnetic Luminex® Assay or the Performance Assay (R&D Systems, Minneapolis, MN), both of which are multiplex assay platforms. These are bead-based systems that use laser sorting of bead populations specific for each analyte, which allows the quantitation of test samples by calculation using standard curves of analytes of known concentrations. Levels of CCL2/MCP-1 and CXCL10/IP-10 were determined by the standard assay system and levels of TNF-α were determined by the Performance Assay. The Performance Luminex® assay has a lower minimum detection range (0.23 to 0.8 pg/ml) than the standard Luminex assay (2 to 90 pg/ml). Each sample was run in duplicate, and the values averaged. All assays were run following the kit instructions, and read on a BioRad BioPlex®100 plate reader, using BioPlex® Manager 6.1 software.

*Cortisol Assay:* Salivary cortisol levels were determined using the Cortisol ELISA Kit (Arbor Assays, Ann Arbor, MI). This is a competitive ELISA for quantitation of cortisol levels. Cortisol levels in subject samples are determined by calculation against a standard curve of known concentrations of the steroid. The assays were carried out according to the kit instructions. Each sample was run in duplicate, and the values averaged. Samples were read at $A_{450}$ using a FLUOstar Omega spectrophotometer (BMG Labtech, Cary, NC). Calculations of cortisol concentrations were determined using PrismGraph 8 (GraphPad Software, San Diego, CA).

*Endotoxin Assay:* Levels of endotoxin in serum were detected by the Pierce® Chromogenic Endotoxin Quantitation Kit (ThermoFisher Scientific, Rockford, IL). This assay is based on the LAL (limulus amoebocyte lysate) reaction, which determines the presence of endotoxin (lipopolysaccharide [LPS]) in solution by the clotting of LAL in the presence of LPS. Levels of endotoxin are determined by calculation against a standard curve of known endotoxin concentrations. This assay can detect levels as low as 0.01 endotoxin units (approx. 100 pg) per ml. Serum was used in this assay since plasma contains anti-coagulant, which would interfere with the LAL reaction. The assay was carried out according to the kit instructions, using only endotoxin-free tubes, plates, and pipet tips. Incubation of the assay was done using a

ThermoFisher digital heating block, set to 37.0° C. Each sample was run in duplicate, and the values averaged. Reading of the chromogenic indicator was done at $A_{405}$ using the FLUOstar Omega spectrophotometer. Calculations of endotoxin levels were determined using Prism Graph 8.

Determination of apremilast concentration in plasma was made by High Performance Liquid Chromatography (HPLC) in the laboratory of Dr. Esther Maier, Drug Dynamics Institute, TherapeUTex, University of Texas – Austin.

*Sample Size:* We used acamprosate as the reference compound for calculating sample size because it is the most recent drug approved for AUD and novel drugs would be expected to exceed its effect size to merit further development. Data from our previous POC study found a moderate effect size for acamprosate (66.1%). This effect size is based on a total VAS craving score of 5.0 ± 1.40 in subjects tested on acamprosate (n=20) and 16.7 ± 3.83 in subjects tested on placebo (n=20), with a coefficient of variation of 100% for each group. Based on these data, 20 completed subjects per treatment group provides adequate power (80%) to detect an effect size of 66.1% at a two-sided significance level of 0.05. Therefore, randomizing 50 subjects (25 per treatment group) allowed for a generous estimate (based on completed studies) of 2-3 noncompliant subjects and 2-3 dropouts for a total of 20 completed subjects per arm. Subjects were block-randomized to conditions using an urn randomization process to ensure balance, as implemented in the R package *randomizr*[20]

**Effects of chronic binge-like drinking on Pde4 gene expression**

HDID-1 female mice (n = 46-48/fluid group) were subjected to chronic binge-like drinking (8-weeks of DID), where mice were offered measured 20% ethanol or water 4-days/week and had access to water (unmeasured) at all other times. 21-hrs after the last DID exposure (when there is no alcohol in the system, reducing possible confounding effects of dose-related issues) in week 8, mice were euthanized by cervical dislocation and rapid decapitation. Whole brains were frozen on powdered dry ice, cryostat sectioned (200 µm), and frozen NAc tissue punches were collected and processed for RNA isolation (using PureZOL and Aurum total RNA fatty and fibrous tissue kit; Bio-Rad). This tissue was generated to determine whether target genes of interest were altered by chronic alcohol drinking. mRNA was reverse transcribed into cDNA using iScript cDNA synthesis kit (Bio-Rad). Quantitative real-time PCR (qPCR) reactions were performed using a CFX384 Touch qPCR system. Reactions were performed in duplicate for each sample to determine relative levels of *18s*, *Pde4a*, and *Pde4b* [Bio-Rad PrimePCR probe assay, targeting mouse genes: Pde4a (5'HEX,3' Iowa Black® FQ, Unique Assay ID:qMmuCIP0030727), Pde4b, 5'TEX 615,3' Iowa Black® FQ, Unique Assay ID:qMmuCIP0036026), and 18s (RPS18, 5' 6-FAM,3' Iowa Black® FQ, Unique Assay ID: qMmuCEP0053856). Data was analyzed using the ddCT method for determining relative gene expression (as described in Falcon et al., 2013 and Ozburn et al., 2015)[21,22].

**Effect of intra-accumbens apremilast administration on binge-like drinking**

*Surgery:* HDID-1 male mice (n = 19-20/fluid group/infusion group; n = 10-11/ethanol group/infusion group for BAL) were administered anesthesia (100 mg/kg ketamine, 10 mg/kg xylazine) and underwent stereotaxic surgery for bilateral guide cannula implantation (coordinates relative to bregma: A/P +1.34, M/L -/+ 0.63, D/V -3.4). The implanted guide cannulae (33-gauge stainless steel, 10 mm long) were aimed at the NAc (core and shell) and were secured by Durelon carboxylate acrylic (3M) and anchored by one stainless steel screw (Small Parts) inserted into the skull. Stainless steel stylets (10 mm, 26 gauge) were inserted into the guide cannula to maintain patency. Mice were provided a nutrient supplement containing carprofen (Medi-Gel) for 3-days prior to surgery (for acclimation and prevention of neophobic response) and 3-5 days post-surgically to aid recovery. Mice received saline, enrofloxacin, and baytril immediately after surgery. Mice were individually housed after surgery to maintain cannula headmount.

*Behavioral testing:* Testing began 7-10 days after surgery. Mice were subjected to 3-weeks of DID testing (4 days/week), where mice were offered 20% ethanol in week 1, water in week 2, and 9.2 mM saccharin in week 3. This procedure was very similar to the above apremilast test, but with two modifications: 1) measured fluid access was limited to 2-hrs on all four days of DID testing and 2) on day 4 of each DID, apremilast was administered 15-20 minutes prior to measured fluid access by direct administration into the NAc. Microinjectors (11 mm length, 26 gauge) were used to deliver 1 µl of vehicle or apremilast (2 µg) at a rate of 0.1 µL/minute for over 10-min. After infusion, microinjectors were left in place for 5-min to allow for diffusion before reinserting stylets. For a subgroup of mice, we collected a 20 µL peri-orbital sinus blood sample for determination of BAL by gas chromatography. After the completion of the study (3 weeks of DID testing), all mice were infused bilaterally with 1 µL of filter sterilized methylene blue dye (17 mg/mL) into the NAc. One hour after infusion, animals were euthanized to extract brains. Whole brains were post-fixed in fresh 4% paraformaldehyde in PBS for 2 days, cryoprotected in PBS + 30% glycerol for 24-hr and sectioned on a freezing microtome (100 µm). Sections were mounted onto slides and inspected using a microscope (localization of methylene blue dye in the NAc was used to confirm injection placement).

**Electrophysiology**

Brain slices were prepared for electrophysiology experiments from adult (13-19 weeks old), male and female (n = 8 male, 6 female), hemizygous *Drd1a*-tdTomato mice, typically within the first few hours of the dark cycle. Mice were anesthetized lightly with isoflurane, decapitated, and their brains were rapidly removed and placed in ice-cold (4°C) high-sucrose ACSF containing the following (in mM): 210 sucrose, 26.2 $NaHCO_3$, 1 $NaH_2PO_4$, 2.5 KCl, 11 dextrose, 6 $MgSO_4$, 2.5 $CaCl_2$, bubbled with 95% $O_2$/5% $CO_2$. Sagittal slices (240 - 250 µm thick) containing the NAc were sectioned in ice-cold, high-sucrose ACSF using a Leica VT1000S vibrating microtome and then transferred to one of two recovery chambers that contained either apremilast (1 µM; final concentration of DMSO = 0.002%) or vehicle (0.002% DMSO) in ACSF (in mM: 124 NaCl, 26 $NaHCO_3$, 1 $NaH_2PO_4$, 4.4 KCl, 10 dextrose, 2.4 $MgSO_4$, 1.8 $CaCl_2$) bubbled with 95% $O_2$/5% $CO_2$. Slices were maintained in these recovery chambers at approximately 32°C for at least one hour prior to transfer to the recording chamber. Recordings were conducted at 30-34°C in ACSF (in mM: 124 NaCl, 26 $NaHCO_3$, 1 $NaH_2PO_4$, 4.4 KCl, 10 dextrose, 1.2 $MgSO_4$, 2 $CaCl_2$) containing either [1 µM] apremilast (final concentration of DMSO = 0.002%) or vehicle (0.002% DMSO), bubbled with 95% $O_2$/5% $CO_{2,}$ and pumped into the recording chamber at ~2.0 mL/min. Recording electrodes (4" thin-wall glass, 1.5 OD/1.12 ID, World Precision Instruments) were made using a P-97 Flaming/Brown micropipette puller (Sutter Instruments) to yield resistances of approximately 4-7 MΩ and contained (in mM): 120 $KMeSO_4$ (ACROS Organics, Geel, Belgium) 12 NaCl, 0.5 EGTA, 10 HEPES, 2 Mg-ATP, 0.3 Tris-GTP (pH 7.3 with KOH, 286 mOsm).

Recordings were acquired on two electrophysiology recording stations, a CV203BU headstage with Axopatch 200B amplifier and a CV-7B headstage with MultiClamp 700B amplifier (Molecular Devices). Experiments for both treatment conditions were performed each day, with the order counterbalanced across days. Approximately half of all recordings were acquired with the technician blind to the treatment condition. Neurons in the NAc core and the border between the shell and core were visually identified using the MRK200 Modular Imaging system from Siskiyou Corporation (Grants Pass, OR) mounted on a vibration isolation table. Cells were identified as D1 MSNs by epifluorescent illumination of tdTomato, while non-fluorescing neurons were selected as putative D2 MSNs on the basis of their morphology. A series of hyperpolarizing and depolarizing intracellular current injections was delivered shortly after obtaining whole-cell configuration and the membrane responses were later used to confirm cells to be MSNs. All cells in the final data set showed membrane voltage responses to this series of current injections that were consistent with features previously reported for MSNs: inward rectification at hyperpolarized potentials, slow ramp depolarization prior to first action potential, and, in the case of repetitive firing, little to no spike frequency adaptation [6,23].

All recordings were filtered at 1 kHz, and digitized at 5 kHz via a Digidata 1440A interface board using Clampex 10.3 (Molecular Devices). Access resistance >30 MΩ or depolarized resting membrane potential (> -60 mV) were criteria for immediate termination of recording.

Excitability was measured by applying depolarizing intracellular current steps (300 msec duration) of increasing amplitude, from 50 pA to 550 pA in 50 pA steps, once every 700 msec, in order to determine rheobase (minimum injected current that elicited an action potential) and the number of action potentials fired at each current amplitude. Input resistance was calculated from the average of the voltage responses to small hyperpolarizing current pulses (-50 pA; 100 ms duration) delivered 100 msec before each 300 msec current step. Action potential characteristics were determined from the first current step to elicit firing, with the exception of the maximum peak to peak frequency, for which the current step eliciting the maximum number of action potentials was used. The action potential threshold was determined as the membrane potential at which dV/dt ≥ 10 mV/msec. AHPs are reported as the difference between the threshold potential and the membrane potential at the respective time point post-action potential: fAHP was determined from the minimum membrane potential within 4 msec of AP threshold; mAHP and sAHP were measured 10 and 15 msec after AP threshold, respectively. All processing of raw data was performed blind to the cell subtype, treatment condition, and sex of the animal.

**Statistics**

All statistical analyses of mouse studies were conducted with GraphPad Prism Software Version 9.0. The sample sizes for each experiment are reported in the appropriate figure legend. If no significant sex X treatment interactions were observed, we performed statistical analyses on data collapsed across sexes. Data are presented as mean +/- SEM.

For behavioral experiments, post-hoc analyses used the Dunnett method to compare all doses to vehicle control group. To ensure that treatment groups did not differ meaningfully before drug treatment, we analyzed data for the first 3 days of each experiment. In experiments where animals were tested in subsequent weeks with a fluid other than ethanol, drug vs. vehicle groups were the same as during the ethanol DID test in week 1, but we analyzed each week's data separately. The principal dependent variables of interest were g/kg ethanol intake and BAL. Intakes for other fluids were analyzed as mL/kg body weight. For stress + CIE and CIE induced escalation in drinking, preliminary analyses of the data indicated that there were not significant variations in intake across the five days of drinking during baseline or each test cycle. Therefore, data were averaged across the last five days of baseline and each test cycle before analysis. Data were analyzed with ANOVA (alpha set at 0.05) followed by Newman-Keuls post-hoc tests whenever a main effect or interaction was significant. Student's t-test was used to analyze data for ethanol intake and BAL data from the intra-accumbens apremilast experiment.

NAc mRNA expression following chronic binge-like ethanol drinking in HDID-1 mice was analyzed as a 1-way ANOVA for fluid type (20% alcohol vs water). Because there were no time effects for either PDE4a and PDE4b NAc gene expression, results were collapsed across time and analyzed as a Student's t-test.

For electrophysiology experiments, membrane properties and action potential characteristics were analyzed using 2-way ANOVA in GraphPad Prism, with cell type (D1 or D2 MSN) and treatment condition (vehicle or apremilast) as between-groups factors. The effect of treatment within each MSN subtype was analyzed using Bonferroni's multiple comparison's test. Action potential firing in response to increasing current steps was analyzed using General Linear

Model Repeated Measures in IBM SPSS Advanced Statistics 23, with current step as the repeated measure, and cell subtype (D1 or D2 MSN) and treatment condition (vehicle or apremilast) as between-groups factors. When sphericity within groups was violated (as indicated by Mauchly's test), the Greenhouse-Geisser adjusted degrees of freedom (rounded to the nearest whole number) and p values are reported in the text.

**Method References**


1. Crabbe, J. C. *et al.* A Line of Mice Selected for High Blood Ethanol Concentrations Shows Drinking in the Dark to Intoxication. *Biol. Psychiatry* (2009) doi:10.1016/j.biopsych.2008.11.002.
2. Crabbe, J. C. *et al.* Progress in a replicated selection for elevated blood ethanol concentrations in HDID mice. *Genes, Brain Behav.* (2014) doi:10.1111/gbb.12105.
3. Anderson, R. I., Lopez, M. F. & Becker, H. C. Stress-induced enhancement of ethanol intake in C57Bl/6J mice with a history of chronic ethanol exposure: Involvement of kappa opioid receptors. *Front. Cell. Neurosci.* (2016) doi:10.3389/fncel.2016.00045.
4. Gruol, D. L., Huitron-Resendiz, S. & Roberts, A. J. Altered brain activity during withdrawal from chronic alcohol is associated with changes in IL-6 signal transduction and GABAergic mechanisms in transgenic mice with increased astrocyte expression of IL-6. *Neuropharmacology* **138**, 32–46 (2018).
5. Gruol, D. L., Melkonian, C., Huitron-Resendiz, S. & Roberts, A. J. Alcohol alters IL-6 signal transduction in the CNS of transgenic mice with increased astrocyte expression of IL-6. *Cell. Mol. Neurobiol.* 1–18 (2020).
6. Ade, K., Wan, Y., Chen, M., Gloss, B. & Calakos, N. An improved BAC transgenic fluorescent reporter line for sensitive and specific identification of striatonigral medium spiny neurons. *Front. Syst. Neurosci.* **5**, 32 (2011).
7. Zarrindast, M.-R., Babapoor-Farrokhran, S., Babapoor-Farrokhran, S. & Rezayof, A. Involvement of opioidergic system of the ventral hippocampus, the nucleus accumbens or the central amygdala in anxiety-related behavior. *Life Sci.* **82**, 1175–1181 (2008).
8. Rhodes, J. S., Best, K., Belknap, J. K., Finn, D. A. & Crabbe, J. C. Evaluation of a simple model of ethanol drinking to intoxication in C57BL/6J mice. *Physiol. Behav.* (2005) doi:10.1016/j.physbeh.2004.10.007.
9. Crabbe, J. C. *et al.* High Drinking in the Dark (HDID) mice are sensitive to the effects of some clinically relevant drugs to reduce binge-like drinking. *Pharmacol. Biochem. Behav.* (2017) doi:10.1016/j.pbb.2017.08.002.
10. Jensen, B. E. *et al.* Ethanol-Related Behaviors in Mouse Lines Selectively Bred for Drinking to Intoxication. *Brain Sci.* **11**, 189 (2021).
11. Sneddon, E. A., Ramsey, O. R., Thomas, A. & Radke, A. K. Increased responding for

alcohol and resistance to aversion in female mice. *Alcohol. Clin. Exp. Res.* **44**, 1400–1409 (2020).

12. Lopez, M. F., Anderson, R. I. & Becker, H. C. Effect of different stressors on voluntary ethanol intake in ethanol-dependent and nondependent C57BL/6J mice. *Alcohol* **51**, 17–23 (2016).

13. Becker, H. C. & Lopez, M. F. Increased ethanol drinking after repeated chronic ethanol exposure and withdrawal experience in C57BL/6 mice. *Alcohol. Clin. Exp. Res.* **28**, 1829–1838 (2004).

14. Finn, D. A. *et al.* Increased drinking during withdrawal from intermittent ethanol exposure is blocked by the CRF receptor antagonist D-Phe-CRF(12-41). *Alcohol. Clin. Exp. Res.* (2007) doi:10.1111/j.1530-0277.2007.00379.x.

15. Griffin III, W. C., Lopez, M. F. & Becker, H. C. Intensity and duration of chronic ethanol exposure is critical for subsequent escalation of voluntary ethanol drinking in mice. *Alcohol. Clin. Exp. Res.* **33**, 1893–1900 (2009).

16. Association, A. P. *Diagnostic and statistical manual of mental disorders (DSM-5®)*. (American Psychiatric Pub, 2013).

17. Sheehan, D. V *et al.* The Mini-International Neuropsychiatric Interview (MINI): the development and validation of a structured diagnostic psychiatric interview for DSM-IV and ICD-10. *J. Clin. Psychiatry* **59**, 22–33 (1998).

18. Sullivan, J. T., Sykora, K., Schneiderman, J., Naranjo, C. A. & Sellers, E. M. Assessment of alcohol withdrawal: the revised clinical institute withdrawal assessment for alcohol scale (CIWA-Ar). *Br. J. Addict.* **84**, 1353–1357 (1989).

19. Sobell, M. B. & Sobell, L. C. Stepped care as a heuristic approach to the treatment of alcohol problems. *J. Consult. Clin. Psychol.* **68**, 573 (2000).

20. Coppock, A., Cooper, J. & Fultz, N. Randomizr: easy-to-use tools for common forms of random assignment and sampling. *R Packag. version 0.20. 0* (2019).

21. Falcon, E., Ozburn, A., Mukherjee, S., Roybal, K. & McClung, C. A. Differential regulation of the period genes in striatal regions following cocaine exposure. *PLoS One* **8**, e66438 (2013).

22. Ozburn, A. R. *et al.* Direct regulation of diurnal drd3 expression and cocaine reward by npas2. *Biol. Psychiatry* (2015) doi:10.1016/j.biopsych.2014.07.030.

23. Ma, Y.-Y. *et al.* Regional and cell-type-specific effects of DAMGO on striatal D1 and D2 dopamine receptor-expressing medium-sized spiny neurons. *ASN Neuro* **4**, AN20110063 (2012).
